# Unveiling the Impact of Arsenic Toxicity on Immune Cells in Atherosclerotic Plaques: Insights from Single-Cell Multi-Omics Profiling

**DOI:** 10.1101/2023.11.23.568429

**Authors:** Kiran Makhani, Xiuhui Yang, France Dierick, Nivetha Subramaniam, Natascha Gagnon, Talin Ebrahimian, Hao Wu, Jun Ding, Koren K. Mann

## Abstract

Millions worldwide are exposed to elevated levels of arsenic. This significantly increases their risk of developing atherosclerosis, a pathology primarily driven by immune cells. While the impact of arsenic on immune cell populations in atherosclerotic plaques has been broadly characterized, cellular heterogeneity is a substantial barrier to in-depth examinations of the cellular dynamics for varying immune cell populations. Here, we present one of the first single-cell multi-omics profiling of atherosclerotic plaques in apolipoprotein E knockout (apoE^-/-^) mice to understand the transcriptomic and epigenetic changes in various immune cells induced by arsenic. Our data reveal that arsenic alters the transcriptional profile of macrophages in a subtype-specific manner with implicated shifts in cell-cell interaction and cell fate predictions. Additionally, our data suggest that arsenic-mediated changes in chromosome accessibility are more profound than their effects on the transcriptome, hence revealing markers of arsenic exposure and potential targets of interventions.

**Teaser:** Arsenic changes gene expression and epigenome primarily of macrophages in atherosclerotic plaque, suggesting intervention targets.

## Introduction

More than 200 million people worldwide are exposed to elevated levels of arsenic, mainly through contaminated food and water (1, 2). Mounting epidemiological evidence implicates arsenic toxicity as a modifiable risk factor for atherosclerosis, the etiology of the majority of cardiovascular diseases (3, 4). Arsenic has been associated with increases in carotid intima-media thickness and plaque score, a measure of atherosclerotic plaque burden over time, in exposed populations (5, 6). Moreover, arsenic exposure is implicated in increased stroke-related hospitalizations and cardiovascular mortalities, both consequences of atherosclerosis (7, 8).

Atherosclerosis is a chronic inflammatory disease of large arteries characterized by fibrous fatty lesions driven primarily by immune cells (9, 10). Broadly, macrophages and dendritic cells in the plaque are known to engulf circulatory lipids, process them and present the antigens to incoming T and B cells (11, 12). T and B cells recognize these antigens and perform their respective effector functions, alongside recruiting more immune cells to the site of the plaque (reviewed in (13)). This crosstalk between plaque-resident immune cells determines their activation states, the cocktail of cytokines they secrete, and, consequently, the progression or regression fates of the plaque. While the diversity of these immune cells and their interactions have been appreciated over the past several decades, the efforts to understand it have been limited by using canonical markers based mostly on in vitro or ex vivo approaches (14). Several studies have evaluated the modulatory effects of arsenic on the individual immune cell types known to constitute an atherosclerotic plaque (reviewed in (15)). For instance, arsenic exposure increases the retention of cholesterol in macrophages in vitro by dampening the reverse cholesterol transport via liver X receptor (LXR) inhibition. Arsenic can result in an altered cytokine profile and oxidative stress in both macrophages and T cells, potential mechanisms of atherogenesis (16, 17). In vivo, arsenic-enhanced atherosclerosis is modelled in apolipoprotein E knockout (apoE^-/-^) mice. While there is an arsenic-induced increase in plaque size and lipid content in this mouse model, the percentage of macrophages in these plaques remains unaltered (18, 19). To understand this, our lab investigated whether arsenic affects the gene expression of the plaque-resident macrophages using in vitro culture of bone-marrow-derived macrophages (20). To this end, we observed that the macrophages polarized with IFN-ɣ or IL-4 responded differently to arsenic exposure (20). However, atherosclerotic plaque is a mesh of different immune cells, and their intercellular interaction makes the plaque microenvironment unique. Therefore, more meaningful insights might be derived from analyzing macrophages resident within the plaque microenvironment.

Single-cell sequencing combines the robustness of next-generation sequencing with the sophistication of microfluidics to isolate individual cells, sequence them, and allow mapping of the transcriptional information to each cell (21, 22). This, along with unbiased dimensionality reduction and clustering algorithms, has fueled the discovery of previously unknown cell subsets at a much higher resolution (21). With the advent of single-cell studies, novel plaque-resident immune cell phenotypes have surfaced and are representative of the consortium of heterogenous cell populations forming the atherosclerotic plaque. ApoE^-/-^ and low-density lipoprotein receptor knockout (ldlR^-/-^) mice are widely used models of atherosclerosis, and single-cell sequencing of the plaques from these mice has resolved the diverse immune cell subsets present in its niche (23, 24). For instance, numerous studies have assigned lesional macrophages to ‘resident-like’, ‘inflammatory’, ‘foamy’, and ‘Trem2^hi^-like’ clusters based on their differential gene expression patterns (25, 26). Importantly, this classification is more elaborate than the classically- and alternatively activated M1 and M2 macrophage subtypes and lies along a spectrum of activation states (27). Additionally, three T cell subsets: the CD8^+^ cytotoxic T cells, CD4^+^ CD8^+^ mixed T cells, and Cxcr6^+^ T cells, three dendritic cell subsets: monocyte-derived, plasmacytoid and mature dendritic cells. B1 and B2 type B cell subsets also have been identified in the atherosclerotic plaques of these mice (24, 25, 28, 29).

Although single cell RNA sequencing (scRNA-Seq) bypasses the limitations of bulk studies, it only profiles the transcriptomes of the cells, hence providing only a partial understanding of the cellular states. Therefore, single-cell multi-omics measurements, capable of profiling the cellular states more comprehensively, are becoming indispensable for single-cell level examinations of varying biological processes, including chronic arsenic exposure. To fill this gap, here, we employ 10x genomics multiome sequencing, which simultaneously profiles the transcriptome and epigenome of the cells of arsenic-exposed apoE^-/-^ mice along with respective controls to ask how arsenic is altering the gene expression profiles of the diverse plaque-resident immune cell subsets to enhance atherosclerosis and to understand the underlying epigenetic mechanisms of these alterations. To our best knowledge, our study is one of the first single-cell multi-omics examinations of arsenic-induced atherosclerosis, which presents unrivaled resources to understand the detrimental effect of arsenic exposure and suggests potential interventions.

Leveraging advanced single-cell technologies, this study delves into the impacts of arsenic exposure on immune cell populations in the atherosclerotic plaque of apoE^-/-^ mice. Our findings resonate with previous studies, revealing the resilience of the cell type diversity within the plaque against arsenic exposure. The macrophages emerged as the dominant immune cell population within the plaque, presenting the most considerable alterations under arsenic exposure. An intriguing pattern emerged when these macrophages were categorized into resident-like and foam cell-like macrophages. Arsenic exposure instigated gene expression changes in these macrophage subtypes, albeit in diametrically opposite directions. As we delved deeper, scRNA-Seq analyses illuminated how arsenic exposure reshapes cell-cell interactions within the atherosclerotic plaque microenvironment, particularly among the macrophage subtypes and other immune cells. The scATAC-Seq data provided a further layer of understanding, uncovering significant epigenetic shifts triggered by arsenic exposure. The macrophage populations identified in this study share considerable overlap with those documented in human atherosclerotic plaques from symptomatic individuals, underscoring the potential clinical and epidemiological relevance of our findings. Thus, this study offers novel insights into the underlying mechanisms of arsenic-enhanced atherosclerosis. Moreover, it underscores the promise of single-cell technologies as powerful tools for toxicology research.

## Methods

### Experimental Design

Our study seeks to comprehensively understand how arsenic affects cellular responses and regulatory networks of immune cells within plaques. Our experimental design integrates multiple techniques, including single-cell RNA sequencing (scRNA-seq) and multi-omics approaches. The primary objectives are to elucidate the effects of arsenic exposure on resident macrophage cells within atherosclerotic plaques and to uncover the molecular mechanisms underlying these effects. Specifically, we aim to identify changes in cell number proportion, gene expression, genomic region accessibility, cell-cell interactions and so on caused by arsenic exposure and find transcription factors as well as regulatory networks based on the changes. The design incorporates components including animal exposure, cell isolation, library preparation, sequencing, and computational analyses to achieve a comprehensive understanding of the molecular responses to arsenic exposure.

### Animal Housing and Exposure

All mice used were approved by the McGill Animal Care and Use Committee. Apolipoprotein E knockout C57BL/6 (Jackson Laboratory, Bar Harbor, Maine) mice were bred in the Lady Davis Institute Animal Facility. Male mice were started on a high fat diet, a low metal purified base diet, (Teklad Custom diet TD.10825: 2016 +15% CB +0.5% Chol, ENVIGO) at weaning and the treatment group was started on 200 ppb arsenic containing water at 5 weeks of age (arsenic was used in the form of sodium arsenite), while the controls were maintained on tap water. For bone marrow-derived macrophages, wild type C57BL/6 male mice between the ages of 8-10 weeks were used. They were bred as mentioned above.

### Plaque CD45^+^ cell isolation

After 13-weeks of exposure, mice were injected with a BB515-conjugated anti-CD45 antibody via tail vein to stain circulating leukocytes and euthanized within the following 10 minutes. Aorta were perfused (intracardiac) with ice-cold PBS. The aortic arch was carefully separated from the perivascular adipose tissue along with the brachiocephalic artery and the carotids, opened and plaque was isolated. Next, the heart was dissected to isolate the plaque from the root of aortic sinus. Both control and treated samples were obtained by pooling the plaques from four mice each for scRNA sequencing. The isolated plaques were digested using a cocktail of enzymes (450 U/ml liberase, 125U/ml collagenase and 60 U/ml hyaluronidase) at 37°C for an hour, then passed through a 70 μm cell strainer to obtain single cell suspensions. The cells were then blocked for 30 minutes at 4°C with anti-mouse CD16/CD32 to block non-specific binding of antibodies to Fc receptors, then stained for CD45^+^ cells, but with an antibody conjugated with BV785 to distinguish plaque-resident leukocytes from those found in the blood. Lastly, 4′,6-diamidino-2-phenylindole (DAPI) was added to exclude dead cells. These plaque resident CD45+ live cells were sorted on a FACS sorter using a 100-um nozzle and sent to the Genome Quebec-Centre for single cell RNA sequencing.

### Library preparation and sequencing

Cell counting was performed manually, with cell viability determined using Trypan Blue and a hemocytometer. Subsequent library construction employed the Chromium Next GEM Single Cell 3 Kit v3.1 (10X Genomics), following the manufacturer’s recommended protocol, with a targeted cell recovery set at 2000 cells. We utilized adapters and PCR primers sourced from 10X Genomics. Library quantification was conducted using the Kapa Illumina GA with Revised Primers-SYBR Fast Universal kit (Kapa Biosystems). Fragment average size was determined using the LabChip GXII instrument (PerkinElmer). In the following steps, libraries were normalized and pooled, followed by denaturation in 0.05N NaOH and neutralization using the HT1 buffer. The resulting library pool was loaded onto an Illumina NovaSeq SP Lane at a concentration of 225pM, employing the Xp protocol as outlined by the manufacturer. The sequencing run was executed for 2x100 cycles in paired-end mode. A PhiX library was incorporated as a control and mixed with the libraries at a 1% level. Base calling was carried out with RTA v3.4.4, after which the bcl2fastq2 v2.20 software was used to demultiplex samples and produce FASTQ reads.

For single cell multiome sequencing, Single-cell suspensions were washed and resuspended in PBS with 0.04% BSA. An aliquot of cells was used for LIVE/DEAD viability testing (Thermo Fisher Scientific). Cells were lysed in lysis buffer (10mM TRIS-HCl, 10mM Nacl, 3mM MgCl2, 0.1% Tween-20, 0.1% NP-40, 0.01% digitonin, 1% BSA, 1mM DTT, 1U/ul RNAse inhibitor) and nuclei isolated according to Nuclei Isolation for Single Cell Multiome ATAC + Gene Expression Sequencing protocol from 10X Genomics (CG000365, Rev C). Afterwards, nuclei were imaged and quantified with DRAQ5 dye (Thermo Fisher Scientific). Single-nuclei libraries were generated using the 10x Genomics Chromium Controller instrument and Chromium Next GEM Single Cell Multiome ATAC + Gene Expression kit (10x Genomics) according to the manufacturer’s protocol (CG000338, Rev E).

The sequencing-ready libraries were cleaned up with SPRIselect Reagent Kit (Beckman Coulter), quality controlled for size distribution and yield (LabChip GX Perkin Elmer) and quantified using qPCR (KAPA Biosystems Library Quantification Kit for Illumina platforms).

Libraries were loaded on independent lanes in an Illumina NovaSeq SP flowcell and sequenced using the following parameters: 51 nt Read1, 10 nt Index1 (i7), 24 nt Index2 (i5) and 151 nt Read2. These cycles were chosen to accommodate both the gene expression library that requires at least 28-10-10-100 and the ATAC library which requires at least 50-8-24-50.

The run demuliplexing was performed using bcl2fastq using a sample sheet. The gene expression lane was demultiplexed using 10-10 index cycles masking the extra bases for Index2. The ATAC lane was demultiplexed with 8bp in single index mode masking the extra bases and exporting the Index2 as a 24bp read containing the ATAC cell barcode. The demultiplexed read files were then processed using "cellranger-arc count" (10X Genomics) which performs the cell barcode and UMI correction, alignment, gene counting and peak calling. Gene-barcode and peak-barcode matrices are then output for downstream analysis.

### FASTQ generation and transcript counting

We undertook the preprocessing of our single-cell RNA-seq data employing the Cell Ranger toolset (v6.1.2). This utility enabled us to demultiplex raw base call (BCL) files, generated via Illumina sequencers, into their FASTQ file counterparts. Concatenation was performed on multiple FASTQ files that originated from the same library and strand, yielding a single unified file. Subsequent read processing, including a suite of operations for raw gene expression quantification, was achieved using the same version of the toolset. This suite entailed read alignment, filtering, barcode counting, and Unique Molecular Identifier (UMI) tallying. Reads were specifically aligned to the mouse reference genome, mm10. Our investigation incorporated Control and Arsenic libraries, effectively integrating them via the aforementioned toolset. The integrated chromium cellular barcodes facilitated the generation of a comprehensive raw cell-by-gene expression matrix, indispensable for all downstream analyses.

### Filtering valid cell barcodes and quality control

The Scanpy library (v1.8.2) (33) operating in a Python (v3.7.0) environment was employed for quality control measures in our single-cell data analysis. Cells that expressed fewer than 200 or more than 2500 genes, or those with more than 10% mitochondrial reads, were deemed of subpar quality and subsequently excluded from further analyses. Post-filtering, the remaining scRNASeq cell-by-gene expression matrix comprised of 2605 cells (1392 cells from the arsenic library and 1213 cells from the control library) and 16636 genes. For the sc-multi-omics RNASeq dataset (scRNASeq (m)), the processed cell-by-gene expression matrix included 5284 cells (2192 cells from the arsenic library and 3092 cells from the control library) and 18050 genes.

### Data normalization and cell dimension reduction

The cell-by-gene expression matrix, post-filtering, was normalized to adjust for library size differences. We carried out this normalization using the Scanpy library (v1.8.2) (33), which was followed by a log transformation. Thereafter, the normalized expression matrix was subjected to a principal component analysis (PCA) to achieve linear dimension reduction. We performed an exploratory analysis on the leading principal components to discern the factors contributing to variance at the cellular level. Upon selection of these key principal components, we established a network of nearest neighbor cells. This network relied on Euclidean distances between cells in the multidimensional PC space, with K=15 neighbors designated for each cell. To finalize, we employed the Uniform Manifold Approximation and Projection (UMAP) method for additional dimension reduction, generating a two-dimensional (2D) representation for improved cell visualization.

### Cell population identification and arsenic enrichment analysis

The cell-by-gene expression matrix was clustered (by cells) to obtain the cell clusters (cell populations) using the Leiden (64) method implemented in Scanpy (33). For each obtained cell cluster, we identified the top expressed genes (potential signature genes) associated with the cluster and annotate the cell type information using the Cellar tool that previously developed, which compared the signature genes associated with each cluster with known cell type markers curated in public databases such as CellMarker (34), PanglaoDB (65), and GSEA (66) to infer the cell identities. The enriched GO terms and pathways associated with the signature genes of each cell cluster were also employed as additional evidence for confident cell type annotation.

In our study, we undertook a series of steps to accurately compare the treated and control groups. We initiated this process by normalizing data for both groups, which allowed us to generate a comprehensible bar graph. Subsequently, we conducted a significance analysis for each cluster, comparing the data between the treated and control groups. In order to determine whether a cluster exhibited significant enrichment in the control or arsenic-treated conditions, we employed a binomial test. This test was utilized to compute an enrichment p-value for every cluster. A cluster was classified as significantly enriched for either control or treated cells if the calculated p-value was less than 0.05. To enhance visual interpretation, we opted to represent these p-values using a logarithmic scale, specifically -log10(p-value). In this graphical representation, a -log10(p-value) greater than 1.3 indicated a p-value less than 0.05, thus marking significant enrichment.

### Differential expression between clusters

One of the notable challenges in scRNA-seq analysis has been the absence of a differential expression model capable of accounting for variability across cellular populations while focusing on differences at the individual subject level, as opposed to treating each cell as an independent sample. Scanpy (v1.8.2) (33) provides a solution to this issue.

Leveraging this tool, we executed a t-test between clusters, which resulted in two comparisons (cluster 4 vs cluster 10 and cluster 3 vs cluster 2). In these assessments, genes exhibiting a log2 fold change either less than -1 or greater than 1, with a p-value of less than 0.05, were classified as significant. We proceeded with a rank rank hypergeometric overlap analysis (67) on the gene lists derived from these two comparisons using the R package RRHO (v1.40.0). Volcano plots were then generated via a free online data analysis website Cloudtutools (http://www.cloudtutu.com). Subsequently, we conducted pathway analyses using KEGG (68) enrichment categories in Webgestalt (69). Pathways displaying a false discovery rate of less than 0.05 were deemed significant (70).

### Cellular trajectory inference

In our study, we harnessed the power of Velocyto (v1.0) (30) within the R (v4.1.2) platform to probe the dynamic landscape of single-cell RNA sequencing data. This step was executed by applying the BAM files from our samples and referring them to the mm10 mouse GTF file to create loom files. To integrate these loom files, we used the loompy (v3.0.7) package in Python (v3.7).

Subsequently, scVelo (v0.2.4) was employed to preprocess individual cells and calculate their momentum. Using the graphical representation of each cell’s momentum generated by scVelo (v0.2.4), we inferred the cellular trajectories between the cell types or clusters of our interest. Importantly, scVelo depicts these trajectories from the perspective of RNA velocity, considering individual cells spliced and unspliced counts.

To complement the velocity-based trajectory inference, we adopted PAGA to delve into the cellular trajectory by assessing the cellular state differences among cells. Scanpy (v1.8.2) (33) was instrumental in this process, aiding in the generation and visualization of edge confidences between each subcluster node across all edges.

In the final stage, we amalgamated the two trajectory inferences using the scdiff2 method to construct an exhaustive trajectory tree with the scdiff2 software. This interactive approach not only allowed us to explore the single-cell data and trajectories but also enhanced our comprehension of the entire process.

### Cell-cell interaction analysis

To elucidate ligand-receptor interaction and cell-cell communication within our scRNA-seq datasets, we utilized CellChat (v1.0.0) (32) in conjunction with R (v4.1.2). The following provides a detailed overview of our approach.

Our initial step involved referencing the Celldb.mouse ligand-receptor database. We used this reference to calculate the communication probability and infer the cellular communication network between cells with the aid of CellChat. This exploration afforded us a nuanced understanding of the cellular crosstalk occurring within our dataset. Post the calculation of communication probabilities, we moved on to aggregate cell-cell communication networks across different clusters. This aggregation facilitated the creation of a holistic view of interactions taking place between distinct cell populations. To visually represent these intricate interactions, we employed CellChat to generate circular plots and heatmaps. These graphical interpretations provided an accessible way to comprehend the complex network of cellular communication within the dataset. Finally, we turned our attention to specific ligand-receptor pairs that spanned across clusters with extensive interaction networks. These key interaction pairs could be visually located in the heatmap presented in Figure 3. By highlighting these significant pairings, we aimed to spotlight potential areas of interest within the complex landscape of cell-cell interactions.

**Figure 3.**
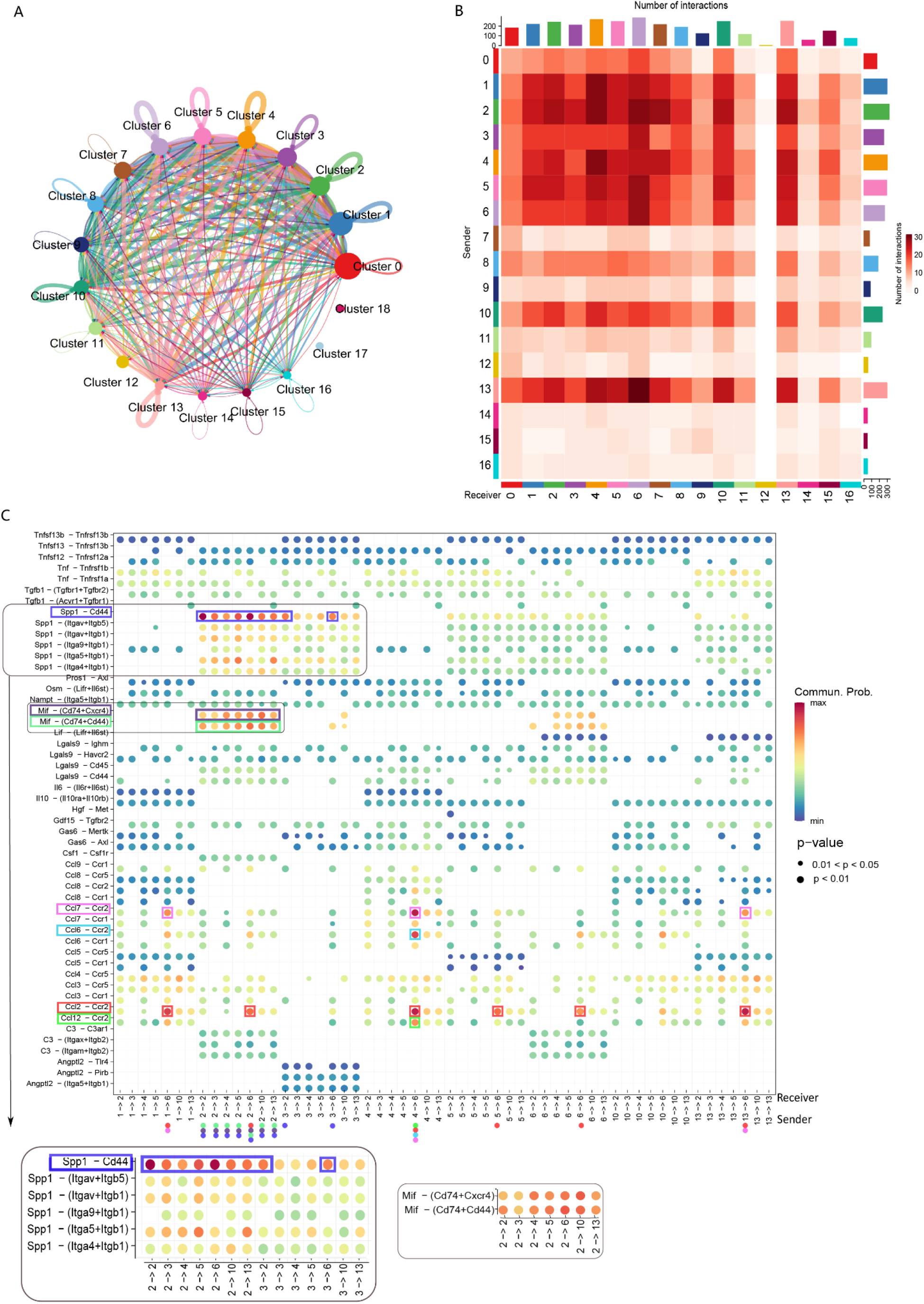
Interactome of immune cells present in an atherosclerotic plaque. Putative cell communication network for the clusters presents in our dataset represented as linkage plot (A) and heatmap (B). In (A), the size of the circles denote the relative abundance of the cluster and the thickness of the loop denote the probability of interaction. In (B), the length of the bars on the top and right side of the plot denote the probability of receiving and sending signals respectively. Comparison of chemokine signals and corresponding receptors represented as dot plot (C). The color of the circle denotes the probability of communication, and the size of the circles denote the p-value of these predictions. Random colors are assigned to the noteworthy interaction pairs for ease of interpretation. Cluster legend: Cluster 0 = Granulocytes | Cluster 1 = Resident-like macrophages | Cluster 2 = Foam cell-like macrophages | Cluster 3 = Foam cell-like macrophages | Cluster 4 = Resident-like macrophages | Cluster 5 = Foam cell-like macrophages | Cluster 6 = THBS1+ macrophages| Cluster 7 = Monocyte-derived dendritic cells/Classical dendritic cells | Cluster 8 = Foam cell-like macrophages | Cluster 9 = Mixed T cells | Cluster 10 = Resident-like macrophages | Cluster 11 = CXCR6+ T cells | Cluster 12 = | Cluster 13 = Foam cell-like macrophages | Cluster 14 = Mature dendritic cells | Cluster 15 = Plasmacytoid dendritic cells | Cluster 16 = B cells | Cluster 17 = Foam cell-like macrophages | Cluster 18 = CD8+ T cells

### Inference of gene regulatory network that dictate the cellular trajectories (Transcription factors)

In our study, we inferred the gene regulatory network through a multifaceted approach, aiming to confidently identify transcription factors and their target genes. We leveraged various sources of information, ensuring our findings were well-supported and robust. Initially, we utilized the scdiff2 method, which allowed us to identify differentially expressed genes between CEC 10 and AEC 4 resident macrophage cells. Alongside these differential genes, we were also able to determine the transcription factors that modulate their expression. This application of scdiff2 provided us with a foundational understanding of the changes between these cell types. Subsequently, we expanded the use of scdiff2 into the scRNA-seq (m) data derived from multiome measurements. This enabled us to identify a set of key transcription factors from another independent perspective, adding an additional layer of verification to our findings.

To further hone our list of key transcription factors driving the shifts of resident macrophage cells, we shifted our focus to epigenomics. We sought transcription factors whose binding sites were enriched in the peaks detected by scATAC-seq with Homer2 (71). This approach allowed us to infer crucial transcription factors from an epigenomic standpoint, which considers the influence of chromatin accessibility on gene regulation. Finally, we integrated the transcription factors identified from the various analyses mentioned above to determine a list of ’confidential’ ones, i.e., those supported by multiple lines of evidence. This integration ensures the reliability of our findings, as transcription factors identified this way are backed by multiple datasets and analytical perspectives. In sum, our approach draws on multiple layers of biological data, marrying transcriptomic, epigenomic, and computational analyses to build a confident picture of the transcriptional regulatory networks in resident macrophage cells.

### Receptor-TF signaling network

In order to comprehend the signaling network that governs the expression differences between control and arsenic-enriched resident macrophages, we harnessed the capabilities of the SDREM (v1.2.0) (38) tool. Our aim was to construct a graph that effectively connects critical cell receptors (identified from cell-cell interaction analysis) with transcription factors inferred using the methods discussed previously. The process started with the input of source nodes (the receptors) and target nodes (transcription factors explaining the expression differences). The SDREM method operates iteratively to identify the transcription factors that not only elucidate the disparities in expression but also best connect the supplied receptors to these target nodes, thereby optimizing network flow.

The SDREM tool comprises two main steps: path-generation and SDREM training. The path-generation step forms the initial network, linking receptors, internal proteins, and transcription factors based on known protein-protein interactions. Then, the SDREM training step refines this network by inferring the most probable signal transduction paths from the receptors to the transcription factors, given the observed differential gene expression patterns.

Following these steps, we obtained a well-defined signaling network that provided an integrated view of the regulatory landscape. To enhance interpretability, we used Cytoscape (v3.9.1) (72), a popular bioinformatics software package for visualizing molecular interaction networks. This allowed us to illustrate the signaling network effectively, offering a tangible representation of the complex regulatory interconnections at play.

In essence, this approach enabled us to trace the route from cell surface interactions to transcriptional changes, thereby providing an inclusive picture of the cellular response to arsenic enrichment in resident macrophages.

### Alignment of the scRNA-seq dataset and the scRNA-seq (m)

In order to align two distinct scRNA-seq datasets, an initial data preprocessing step was performed as outlined above. Following this, we utilized the Scanpy function (v1.8.2) (46) "rank_genes_groups" to obtain the top 100 differentially expressed genes. These genes were then leveraged to evaluate the correspondence between clusters in the scRNA-seq dataset and those in the scRNA-seq (m) dataset.

To quantitatively test the correlation between each pair of clusters across these two datasets, we used the SuperExactTest tool (https://network.shinyapps.io/superexacttest/) (85). This tool applies a hypergeometric test, a statistical method that can accurately measure the overlap between two gene sets. This process enabled us to identify the most highly correlated cluster pairs across the two datasets, which were subsequently aligned.

However, in instances where no significant pair correlation was observed for a particular cluster, we chose not to enforce alignment. This approach of stringent alignment based on robust statistical measures ensured that only highly similar cell populations across the two datasets were mapped onto each other, thereby maintaining the integrity of the biological insights derived from subsequent analyses.

### Single cell multi-omics ATAC data sequencing analysis workflow

Our approach to infer regulatory networks from ATAC-seq data began with obtaining the samples, as described in the previous section. The data was preprocessed using cisTopic (v2.1.0) (48), which enabled us to summarize peak data into a manageable number of topics. Specifically, we derived 100 features or ’topics’ from the fragment files, bed files, and matrix files of the datasets.

To correct for batch effects and achieve data integration, we used Scanorama (v1.7.2) (86). We then applied UMAP for dimensionality reduction, followed by the Leiden algorithm (77) for cell clustering. We transferred the cluster labels from scRNA-seq(m) data to scATAC-seq data utilizing Scanpy (v1.8.2) (46). This process resulted in an ATAC-seq dataset consisting of 7547 cells (2940 from the Arsenic library and 4607 from the control library), represented by 100 topics.

To visualize the cell-topic distributions for each cluster, we employed cisTopic (v2.1.0) (48) again. We transformed the most expressed topic of a cluster into a gene list using PAVIS2 (https://manticore.niehs.nih.gov/pavis2/) (76), based on the peaks contained within the topic. These derived genes enabled various downstream analyses.

For instance, we conducted pathway analysis for clusters 5 and 6 (labels transferred from the scRNA-seq (m)) using the ToppGene website (https://toppgene.cchmc.org/enrichment.jsp) (77). Moreover, the transcription factor enrichment analysis for the scATAC-seq data, discussed in the above section, was also performed with these derived genes. This multi-steps, multi-tools approach allowed us to infer detailed and reliable regulatory networks from the ATAC-seq data.

### Transcriptome and epigenome visualization

In our study, we visualized genomic data derived from our multi-omics datasets utilizing a series of bioinformatics tools. Initially, we processed the Binary Alignment Map (BAM) files from our multi-omics datasets. For this, we employed the split function provided by Bamtools (v2.5.1) (75) to divide the BAM files into manageable segments. Following this, we utilized the bamCoverage function in Deeptools (v1.5.12) (78). This tool allowed us to normalize the BAM files and convert them into BigWig format, a binary format designed for dense, continuous data and preferred for its ease of visualization. Lastly, we employed the Integrative Genomics Viewer (https://igv.org/) (79), a high-performance genomics data visualization tool, to visualize the converted BigWig files. We referenced the mouse genome (mm10) during the visualization process. Through this series of steps, we were able to create accessible and detailed genomic visualizations, enabling further insights into the data collected from our multi-omics datasets.

### Human data correlation

In an effort to validate our findings in human samples, we employed a human single-cell dataset (39) for comparative analysis. We initially preprocessed the human data utilizing the pipeline described above, ensuring compatibility and comparability with our mouse datasets. Dimensionality reduction via UMAP and the identification of differentially expressed genes were performed using Scanpy (v1.8.2) (33). These steps enabled us to gain an overview of the human cellular landscape and identify genes driving differences between cell populations. Next, we employed the SuperExactTest tool (https://network.shinyapps.io/superexacttest/) (73) to perform a hypergeometric test, examining the correlation between each pair of clusters in the human and mouse datasets. This provided a statistical measure of the overlap between gene sets in the two species, supporting the comparative analysis.

In order to accurately align human and mouse gene identifiers, we utilized the NCBI HomoloGene database (https://www.ncbi.nlm.nih.gov/homologene) (80). This tool facilitated the conversion of human gene symbols to their mouse counterparts, ensuring the accuracy of our comparative analyses and supporting the translation of our mouse model findings to human biology.

### Statistical analysis

No statistical methods were used to predetermine sample size. Experiments were not randomized, and investigators were not blinded to allocation during library preparation, experiments, or analysis. P-values in our study were all adjusted with false discovery rate (FDR). A P-value of <0.05 was considered statistically significant. No prior knowledge was used in data preprocessing.

## Results

### scRNA-Seq delineates immune cell populations in an atherosclerotic plaque

Arsenic exposure is associated with enhanced atherosclerosis in apoE^-/-^ mice that is accompanied by changes in the plaque constituents: decreased collagen and smooth muscle cell staining and increased lipid retention (18, 19). Notably, macrophage staining was not altered, suggesting that arsenic may impact macrophage phenotype rather than quantity (19). To test this hypothesis, we utilized scRNA-Seq to examine the diversity of plaque-resident immune cells in apoE^-/-^ mice comparing control and arsenic-treatment. To do so, 5-week-old male apoE^-/-^ mice were fed a high-fat diet and maintained on either tap water or 200 ppb arsenic-containing water for 13 weeks. Atherosclerotic plaques were enzymatically digested, and live plaque-resident CD45^+^ cells were isolated using FACS and used for scRNA-Seq (Figures 1A and S1). Post quality control, 1211 cells from control mice and 1392 cells from arsenic-exposed mice were analyzed. With our scRNA-Seq data analysis, we identified 19 distinct cell clusters based on gene expression patterns (Figures 1B, S2A and Table S1). Monocyte/macrophage, dendritic cells, and T cells constituted the majority of CD45^+^ cells. Macrophages, the most abundant cell type, constituted 62.7% of the total cells. Clusters with high expression of *Cd3e*, indicative of T cells/NKT cells, accounted for 10.3% of the dataset. B cells and dendritic cells made up 0.6% and 9.2% of total CD45^+^ cells, respectively. Among the dendritic cells, cluster 7, the largest group, displayed high expression of *Cd209*, *Ifitm1*, *Ifi30*, and *Klrd1*, suggesting monocyte-derived or classical dendritic cells. Cluster 14, marked by high *Ccr7* and *Fscn1* expression, pointed towards mature dendritic cells, while cluster 15, expressing high levels of *Ifi205*, *DNase1l3*, and *Siglech*, was associated with plasmacytoid dendritic cells.

**Figure 1.**
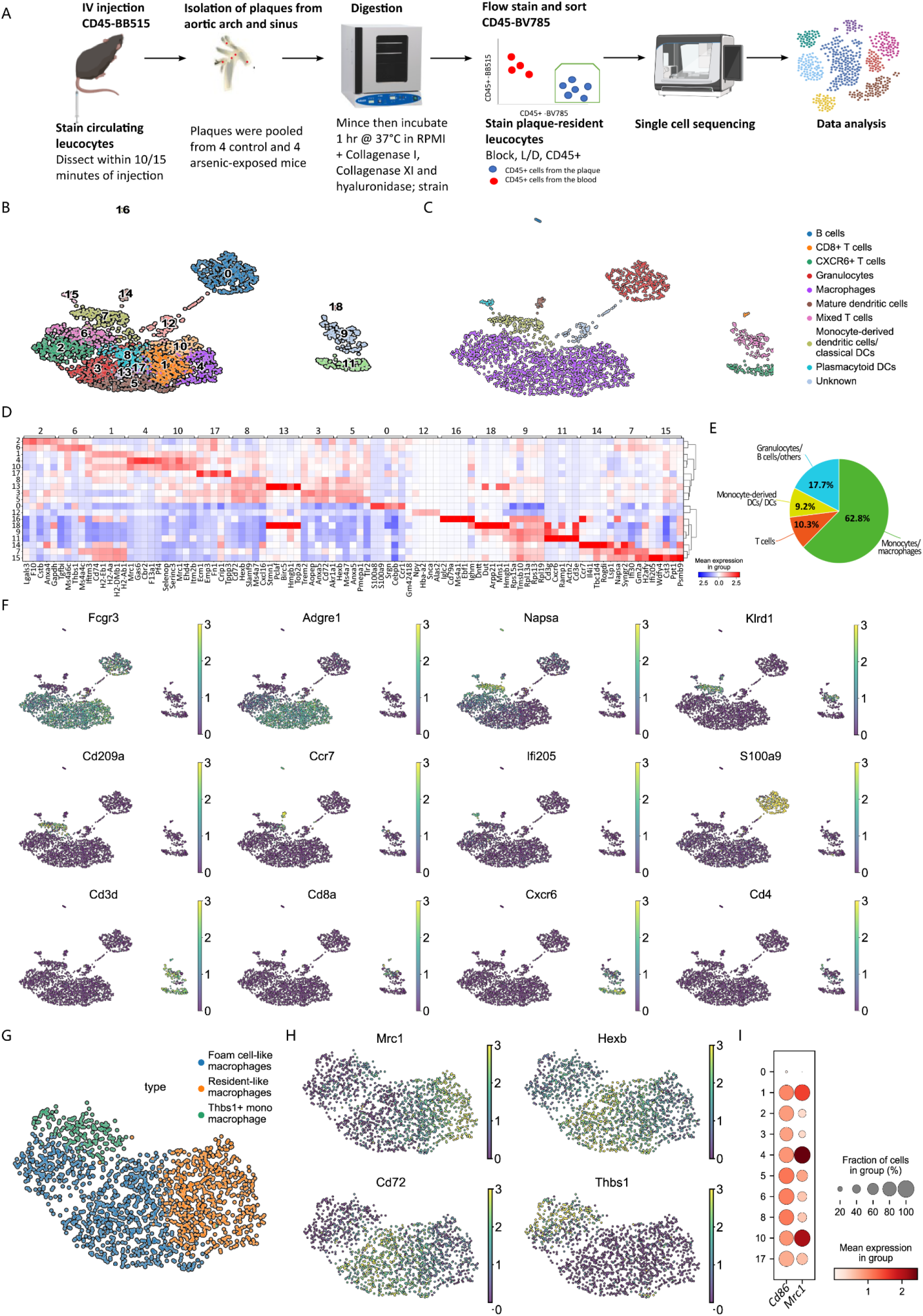
Characterisation of immune cell populations in an atherosclerotic plaque. Schematic of the experimental design (A). Uniform Manifold Approximation and Projection (UMAP) of the gene expression data in single cells extracted from control and arsenic-exposed atherosclerotic plaques of apolipoprotein E knockout (apoE^-/-^) mice segregated into 19 clusters using unsupervised clustering (B) and annotated for the immune cell types (C). Heatmap showing the 5 most upregulated genes in each cluster defined in ‘C’ (D). Relative abundance of the different immune cell types in the integrated dataset (E). Gene expression patterns of Fcgr3, Agre1, Napsa, Klrd1, Cd209a, Ccr7, Ifi205, S100a8, Cd3d, Cd8a, Cxcr6, Cd4 to represent the cell types identified in ‘C’ (F). Macrophage subtypes present in the integrated dataset represented by gene expression data of all cells (G) as well as gene expression patterns of Mrc1, Hexb, Cd72, Thbs1 (H) on a UMAP. Evaluating pro-inflammatory (M1) and pro-resolving (M2) macrophage phenotypes in our dataset, characterized by Cd86 and Mrc1 gene expression respectively (I).

Due to their prevalence and diversity, we further delineated macrophage subtypes (Figures 1C-F, S2 and Table S2). Based on previous scRNA-Seq datasets, at least three different types of macrophages exist within murine plaques (24). Resident macrophages, found in clusters 1, 4, and 10, expressed higher levels of *F13a1*, *Lyve1*, and *Mrc1*. Clusters 2, 3, 5, 8, 13, and 17 highly expressed classical markers of foam-like macrophages (Figure 1G and 1H). Intriguingly, macrophage cluster 6 highly expressed Thbs1 and did not align with the known macrophage subtypes, thus were labelled Thbs1^+^ macrophages. However, this cluster seemed to be a mix of monocytes and macrophages (24, 25) (S2). Notably, none of the macrophage clusters could be classified as pro-inflammatory (M1; marked by *Cd86* expression) or pro-resolving (M2; marked by *Mrc1* expression), in line with other single cell studies in mice (Figure 1I).

### Arsenic exposure changes the relative composition of macrophages

We next segregated data and annotated cells based on treatment group, control versus arsenic-exposed, in order to identify arsenic-induced alterations in the abundance of plaque-resident immune cells and their subtypes after normalization for total cell count (Figure 2A). The subsequent analysis was aimed at identifying any significant alterations in the abundance of plaque-resident immune cells and their subtypes within the plaques of arsenic-exposed mice, normalized for total cell count. Remarkably, despite arsenic exposure, the overall proportion of macrophages, T cells, and dendritic cells, among other immune cell populations, remained unaltered within the plaques (Figure 2B). However, there were notable differences between clusters that were enriched in either control or arsenic-exposed mice. Specifically, clusters 0, 2, 5, 6, and 10, henceforth termed as <u>C</u>ontrol <u>E</u>nriched <u>C</u>lusters (CECs), were significantly enriched in cells from control samples, while clusters 1, 3, 4, 8, 12, 13, and 17, henceforth referred to as <u>A</u>rsenic <u>E</u>nriched <u>C</u>lusters (AECs), were enriched in cells from arsenic-exposed mice (p-value: <0.05) (Figure 2C and Table S3). Interestingly, nearly all the differentially enriched clusters were of monocyte/macrophage origin, with no significant enrichment observed in dendritic cell, T cell, or B cell clusters. Upon aggregating the macrophage cells from both conditions, we discovered a significant increase in resident-like macrophages in arsenic-treated samples (control: 221 and arsenic-treated: 370; p-value: 2.69x10^-8^). In contrast, the changes in foam cell-like macrophages were statistically non-significant (control: 399 and arsenic-treated: 462; p-value: 0.514). Thus, arsenic has little effect on the percentage of cell types within the plaque, with the exception of resident-like macrophages. However, arsenic significantly alters the composition of macrophage populations.

**Figure 2.**
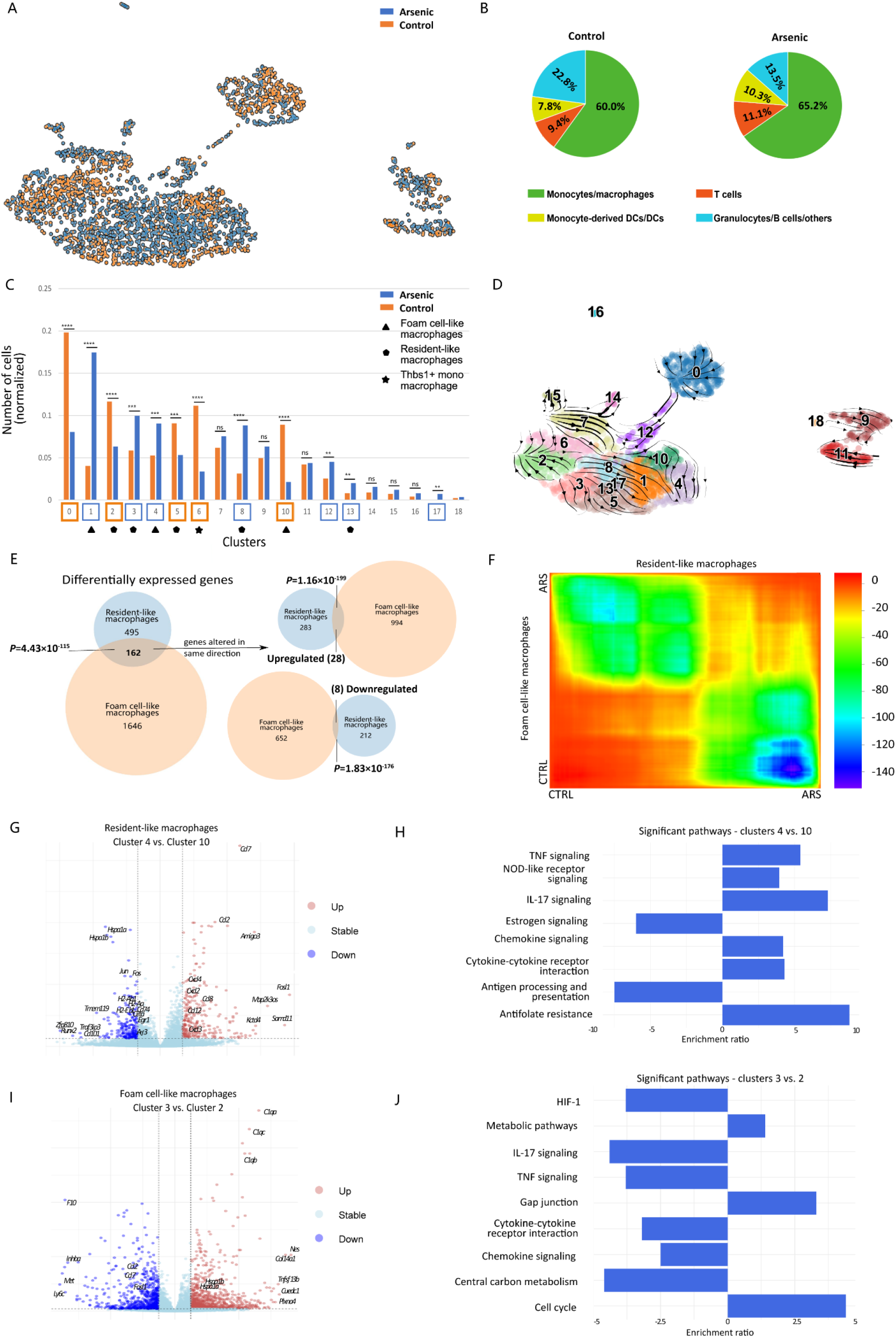
Exploration of diverse macrophage populations. Two dimensional UMAP of the gene expression data in single cells extracted from control (orange) and arsenic-exposed (blue) aorta of apolipoprotein E knockout (apoE^-/-^) mice (A). Relative abundance of different immune cell clusters in control (left) versus arsenic-treated (right) samples as normalized number of cells (B) and as percentage of total CD45^+^ cells (C). RNA velocity plot predicting transcriptomic changes of the different cell clusters; the length of the arrows represents the transcription rate (velocity) and the directionality is towards the inferred future state (D). Venn diagram of the differentially expressed genes overlapping between resident-like (clusters 4 as compared to cluster 10) and foam cell-like (cluster 3 as compared to cluster 2) macrophages (E). Heatmap showing the global gene expression changes in the resident-like and foam cell-like clusters in the cells obtained from control versus arsenic exposed atherosclerotic plaques (F). Volcano plots showing the differentially expressed genes (G) and bar plots representing differentially altered pathways (H) between resident-like clusters 4 and 10 as a result of arsenic exposure; significantly upregulated genes (log2FC > 1; p < 0.005) are in red and significantly downregulated genes (log2FC > -1; p < 0.005) are in blue. Volcano plots showing the differentially expressed genes (I) and bar plots representing differentially altered pathways (J) between foam cell-like clusters 3 and 2 as a result of arsenic exposure; significantly upregulated genes (log2FC > 1; p < 0.005) are in red and significantly downregulated genes (log2FC > -1; p<0.005) are in blue.

### Trajectory inference analysis suggests the potential cellular dynamics associated with macrophages under arsenic

Pseudo-time analyses help determine the fate of cell populations and infer their trajectories of differentiation in single cell datasets. It utilizes the relative abundance of nascent and mature mRNAs to estimate the rate of splicing in order to predict directionality (30). Therefore, to estimate the gene expression changes in the diverse single cell clusters identified in our study, we performed RNA velocity analysis, a pseudo-time approach (Figure 2D), and focused on the macrophage clusters. Amongst resident-like macrophages, CEC 10 likely transitions into AEC 4 upon arsenic exposure (Figure 2D), suggesting that arsenic exposure shifts the resident macrophage population, but not toward foam cell-like phenotypes. Amongst the larger foam cell clusters, CEC 2 maintains gene expression homeostasis, whereas AEC 3 has plausible trajectories towards several other smaller foam cell clusters 5 (CEC), 8 (AEC), 13 (AEC) and 17 (AEC), before terminating into AEC 1 (Figure 2D). While we did not find any inflammatory macrophage clusters in our dataset, clusters 5, 8, 13 and 17 have relatively higher expression of genes (*Ccl2*, *Ccl3*, *Ccl4*, *Cxcl2*) reminiscent of these macrophages. Our data indicate that arsenic treatment increases the heterogeneity of the foam-like cells within the plaque.

Moreover, we observed that the Thbs1^+^ monocytes/macrophage CEC 6 is one of the most dynamic clusters and has probable trajectories terminating in foam cell-like CEC 2 and resident-like AEC 1 (Figure 2D). Studies have shown that newly recruited monocytes/macrophages can attain both resident-like and foam-like phenotypes (31). Therefore, our observations are consistent with previous findings and indicate that these transitions are part of the normal pathogenesis in an apoE^-/-^ mouse.

Overall, our data suggest that instead of coalescing different macrophage subtypes, arsenic changes the foam- and resident-like macrophages in unique ways. While arsenic appears to shift the resident-like macrophage cluster into a single arsenic-enriched, resident-like subtype, arsenic increases the heterogeneity of foam cell-like macrophages.

### Arsenic induces distinct gene expression changes in resident-like and foam cell-like macrophages

We next turned our attention to the comparison of arsenic-induced gene expression changes within the two dominant macrophage types, the resident-like and foam cell-like clusters. The impact of arsenic was markedly more significant on foam cell-like macrophages, altering 1,808 genes, in comparison to 657 genes in resident-like macrophages (Figure 2E), under the same cutoffs. Interestingly, only 36 genes experienced alterations in the same direction (28 upregulated, 8 downregulated) in both cell types due to arsenic exposure (Figure 2E). The majority (∼78%) of the shared genes (126 of 162) were modified in opposite directions (Figure 2E). This lack of overlap and discrepancy in the direction of the change between resident-like and foam cell-like differential gene expression, as depicted by a selection of such genes in Figure 2F, prompted us to examine each macrophage type individually.

To illuminate the drivers of the observed transitions revealed through RNA velocity analysis, we compared the differentially expressed genes between resident-like CEC 10 and AEC 4. We identified 495 significantly altered genes, with 283 upregulated and 212 downregulated (Figure 2G). Pathway analysis identified an upregulation of cytokine/chemokine signaling and decreased antigen presentation in AEC 4 (Figure 2H and Tables S4, S6-S7). In contrast, when we compared the foam-cell like CEC 2 and AEC 3, 1646 genes were significantly different (994 upregulated; 652 downregulated; Figure 2I). Pathways modulated by arsenic included cell cycle, gap junctions, HIF signaling, and central carbon metabolism (Figure 2J and Tables S5, S8-S9). Of note, cytokine/chemokine signaling and antigen presentation were also identified as pathways modulated by arsenic, however, in contrast to resident-like macrophages, these pathways were decreased in foam-cell macrophages. These findings underscore the distinct ways in which arsenic alters gene expression patterns in different macrophage subtypes and suggest that arsenic utilizes unique pathways to mediate pro-atherosclerotic changes in these distinct macrophage populations.

### Arsenic exposure substantially shifts cell-cell interaction patterns associated with the macrophages

Hypothesizing that unique alterations in distinct macrophage subtypes stem from specific intercellular interactions, we explored dynamic cell-cell communication within the plaque microenvironment as this could potentially influence atherosclerosis progression or regression. Employing CellChat (32), a tool predicting communication likelihood between cells based on ligands, receptors, and co-factors, we assessed interactions across cell clusters. Figure 3A portrays predicted ligand-receptor interactions, where the circle size reflects the abundance of cells in each cluster and edge width indicates communication probability. Figure 3B depicts a heatmap of interaction quantities between communicating clusters, x- and y-axes denoting receivers and senders, respectively. Our single-cell dataset unveiled 3064 interaction pairs, with clusters 1-6, 10, and 13 being prominent (Figures 3A-B). Specifically, clusters 6 and 4 seemed to receive the majority of signals (290 and 266 pairs separately, top two in all clusters), while clusters 2 and 4 were the prominent signal senders (329 and 301 pairs separately, also top two in all clusters) (Figure 3A-B). Notably, the majority of active cell-cell engagement clusters comprised of monocytes and macrophages (95% of pairs).

Resident-like macrophages displayed elevated chemokines post-arsenic exposure (Figure 2G-H). This was reflected in the CellChat interactome with chemokine-receptor pairs (Figure 3C). We studied the changes in ligand-receptor interactions due to arsenic exposure. Arsenic exposure led to increased interactions in resident-like clusters 4 and 10 and decreased interactions in foam-like clusters 2, 3 and 5 (Figure 3A-B). For example, AEC 4 showed elevated interactions with incoming monocytes/macrophages (cluster 6) and foam-cells, reflective of chemokine-receptor upregulation, such as that of CCL2/CCL7/CCL12-CCR2 and CCL6-CCR2, consistent with expression data (Figure 2G and 3C). In contrast, foam-cell cluster 2 and 3 (AEC) showed reduced interactions, especially with cluster 6 (Figure 3B), aligning with decreased chemokine expression (Figure 2I-J). Interestingly, AEC 2 and 3 had heightened interactions with resident-like clusters, suggesting potential foam-cell involvement in recruiting and shaping resident-like macrophages (Figure 3B).

Cluster 6, a mix of monocytes and macrophages, notably received recruitment and migratory signaling via CCL2-CCR2 and CCL7-CCR2 ligand-receptor pairs due to high Ccr2 expression. Cluster 6 also exhibited autocrine signaling along the CCL2-CCR2 axis. Unique to resident-like AEC 4, it was predicted to relay chemotactic and migratory signals via CCL6/CCL12-CCR2, facilitating transition of newly recruited monocytes/macrophages into resident-like phenotypes. These data suggest that arsenic changes the migratory signals, thus the communication between the macrophage subtypes.

Foam cells, known for their high SPP1 expression in atherosclerotic plaques, primarily mediate crosstalk with other monocyte/macrophage clusters via CD44 and integrins. In our study, the SPP1-CD44 interaction dominated foam cell-driven cell-cell communication preferentially over integrin receptors (Figure 3C). Intriguingly, the SPP1-CD44 interaction was highest within the CEC 2 cluster as compared to AEC 3. These data suggest that arsenic may alter the receptor engagement by SPP1 within plaque macrophage populations and point to the role of arsenic in modulating the interactions of foam cells with other cell types via SPP1-CD44, which may have important implications for the progression of atherosclerosis.

The MIF-CD74+CXCR4/CD74+CD44 cytokine interactions were unique to the foam-like CEC 2 and thus, were absent in the foam-cell AEC 3 (Figure 3C). This interaction was primarily with the resident-like clusters, suggesting an arsenic-induced change in the foam cell-resident macrophage crosstalk. However, the biological implications of this change and its relevance to atherosclerosis progression are yet to be fully elucidated.

Our single-cell ligand-receptor interactome provides a detailed view of the dynamic and complex cell-cell communication within the plaque microenvironment. The changes in intercellular communication, possibly driven by arsenic exposure, could underlie the distinct alterations observed in resident-like and foam-like macrophages, and provide novel insights into the mechanisms of arsenic-induced atherosclerosis.

### The single-cell mutli-omics data confirms and enhances scRNA-seq findings

In the preceding sections, we delineated the various cell types identified in our scRNA-seq dataset and scrutinized the arsenic-induced shifts in macrophage subpopulations – specifically, resident-like and foam cell-like macrophages. To further probe the epigenetic adjustments accompanying these transcriptional alterations, we conducted single-cell 10x Genomics multiome sequencing, employing a new set of animals for plaque isolation and performing paired single-cell RNA and ATAC sequencing. This allows for a comprehensive examination of transcriptome and epigenome of the immune cell populations in plaques.

We began by integrating data from both control and arsenic-exposed mice derived from the RNA-seq component of the multiome (denoted henceforth as scRNA-Seq (m)), and established cell clusters based on the gene expression patterns discerned earlier. Scanpy-assisted clustering yielded 20 unique cell clusters (Figures 4A, S3A and Table S10), presented on the UMAP in Figure 4A alongside their corresponding scRNA-Seq clusters in parentheses (33). We then attributed biological identities to these clusters using CellMarker and the murine gene expression atlas (34) (Figures 4B and S3B). Consistent with the initial scRNA-seq, the multiome scRNA-Seq (m) discerned foam- and resident-like macrophages, granulocytes, B cells, CD4^+^ and CD8^+^ T cells, and dendritic cells. Interestingly, an NKT cell cluster, absent from our initial dataset, was also discerned, potentially due to the larger number of profiled cells (a total of 5284 cells (Table S10) compared with scRNA-seq’s 2603 cells (Table S1)). Representative gene signatures of each immune cell subtype, utilized for annotation, are illustrated in Figures 4C, S3 and Table S11. Clusters 0-3 were classified as foam-like macrophages, while clusters 5-7, 9, 11, and 16 exhibited gene signatures of resident-like macrophages. Among non-macrophage clusters, clusters 4, 14, and 17 were identified as DCs/moDCs, clusters 8, 12, 13, 18, and 19 as T cells, cluster 10 as granulocytes, and cluster 15 as a B cell cluster (Figure 4B). To provide a comparative view of the cluster characteristics between scRNA-Seq and scRNA-Seq (m), we enumerated the cells attributed to the various immune cell types (Figure 4D). Excluding granulocytes, the relative ratios of the annotated immune cells were generally consistent between the two datasets.

**Figure 4.**
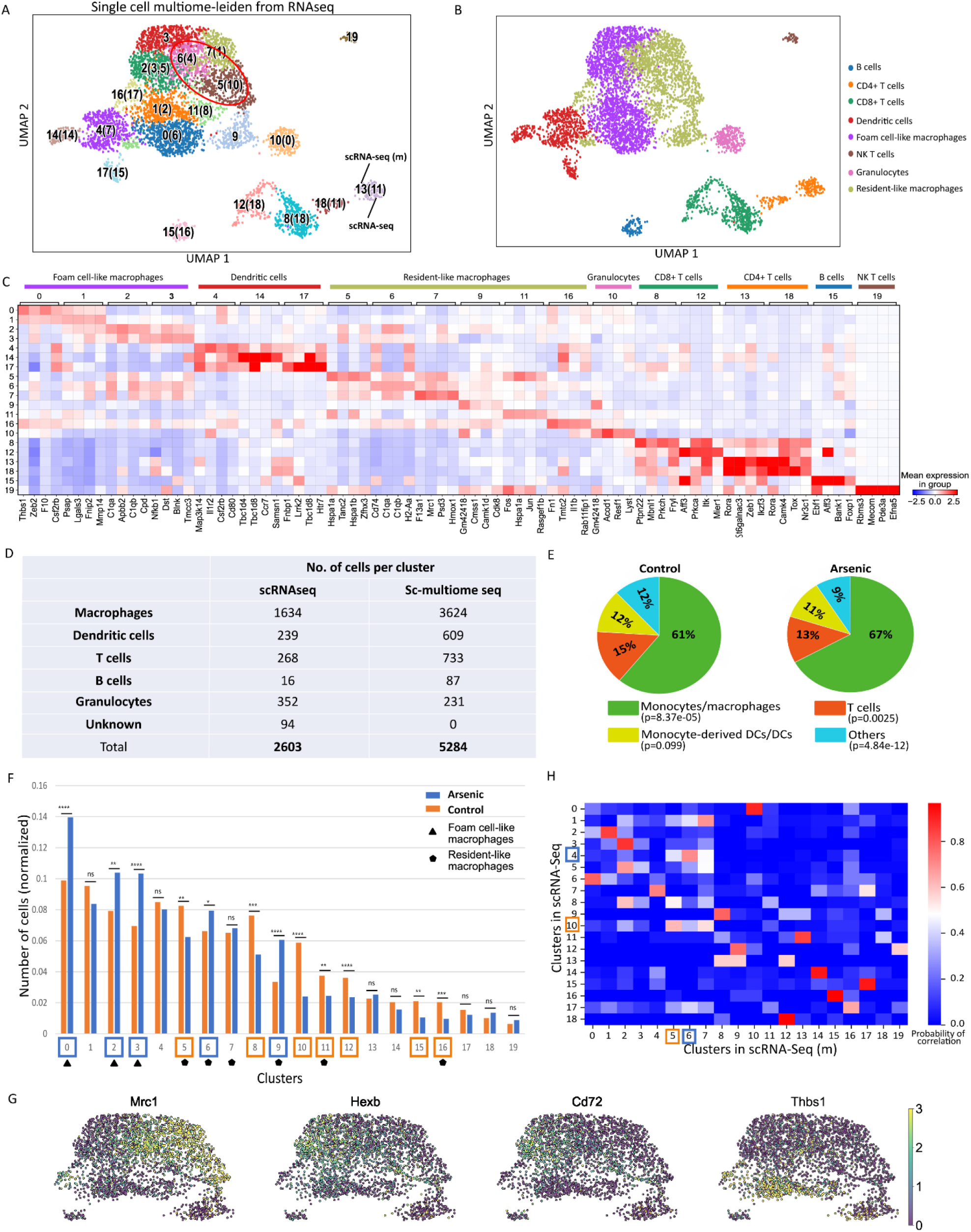
Characterization of the immune cell landscape from scRNA-Seq(m) dataset. Uniform Manifold Approximation and Projection (UMAP) of the single cell gene expression data extracted from scRNA-Seq(m) performed on control and arsenic-exposed atherosclerotic plaques of apolipoprotein E knockout (apoE^-/-^) mice segregated into clusters using unsupervised clustering, where the numbers in parantheses represent the corresponding clusters in the scRNA-Seq dataset (A) and annotated for the immune cell types (B). Heatmap showing the top most upregulated genes in each cluster defined in ‘B’ (C). Comparison of the number of cells pertaining to each of the immune cell types deduced from scRNA-Seq and scRNA-Seq(m) (D). Pie charts presenting the relative abundance of different immune cell clusters in control (left) versus arsenic-treated (right) samples as a percentage of total CD45^+^ cells (E) and as normalized number of cells (F). Macrophage subtypes present in the scRNA-Seq(m) dataset as delineated by the gene expression patterns of Mrc1, Hexb, Cd72, Thbs1 on a UMAP (G). Correlation plot comparing the immune cell clusters in scRNA-Seq and that in scRNA-Seq(m).

Subsequently, we quantified the relative abundance of each immune cell cluster in control and arsenic-treated samples from scRNA-Seq (m). In both groups, the majority of CD45^+^ cells were comprised of monocytes/macrophages, followed by dendritic cells and T cell clusters (Figure 4E). We then examined the arsenic-induced effects on the relative abundance of the identified cell clusters. Clusters 0, 2, 3, 6, and 9 were predominantly enriched in cells from treated animals, whereas clusters 5, 8, 10, 11, 12, 15, and 16 were more prevalent in cells from control animals (Figure 4F and Table S12).

Our focus then turned to macrophage subtypes analogous to those exhibiting intriguing patterns in our scRNA-Seq dataset. We annotated macrophage subtypes in the new dataset using canonical markers from the previous dataset (Figure 4G), identifying markers of foam- and resident-like macrophages, as well as *Thbs1*+ macrophages. We also discovered resident-like clusters 5 and 6 in the scRNA-Seq (m) dataset analogous to scRNA-Seq CEC 10 and AEC 4, respectively (Figures 4H, S2A, S3A and Tables S2, S11). The substantial overlap between the two scRNA-Seq data sets confirmed the reproducibility of our sample processing and analyses. However, despite the sophistication of single-cell transcriptomics, deducing regulatory code remains challenging due to gene regulation being orchestrated by a myriad of transcription factors binding to enhancers and promoters. To overcome this, we employed the scATAC-Seq data from the multiome sequencing dataset to understand chromatin accessibility profiles linked to the impending transcriptional changes. The control and arsenic scATAC-Seq data were first integrated to cluster single cells based on differential chromatin accessibility using cisTopic (35). This unsupervised Bayesian framework groups genomic regions into regulatory topics and clusters cells based on their regulatory topic contributions (35). As a result, the single cells were grouped into 19 distinct clusters with signature topics representing each of these clusters (Figures 5A and 5C). Here, the term scATAC-seq "topic" inferred by cistopic (35) refers to a collection of co-accessible regions (or peaks) across the genome, similar to the concept of "topics" in document classification tasks. In the context of scATAC-seq data, a "topic" is essentially a set of genomic regions that tend to be open (i.e., accessible to the transposase enzyme) together across cells, which may suggest co-regulation by the same set of transcription factors. The scRNA-Seq (m) data was then used to assign identities to the scATAC-Seq clusters (Figure 5B). All the clusters from the former were recapitulated in the latter, with three unidentified clusters that did not correspond to any in the scRNA-Seq, hence labeled ’NA’ in Figure 5B. Post the preprocessing of our single-cell ATAC-seq data, we observed no substantial batch effects. This outcome was corroborated through the application of batch effect removal methods like Harmony (36) and Seurat (37). Notably, the Uniform Manifold Approximation and Projection (UMAP) maintained consistency even after the application of these batch effect removal tools. For a thorough overview of the process, we direct you to Figure S4.

**Figure 5.**
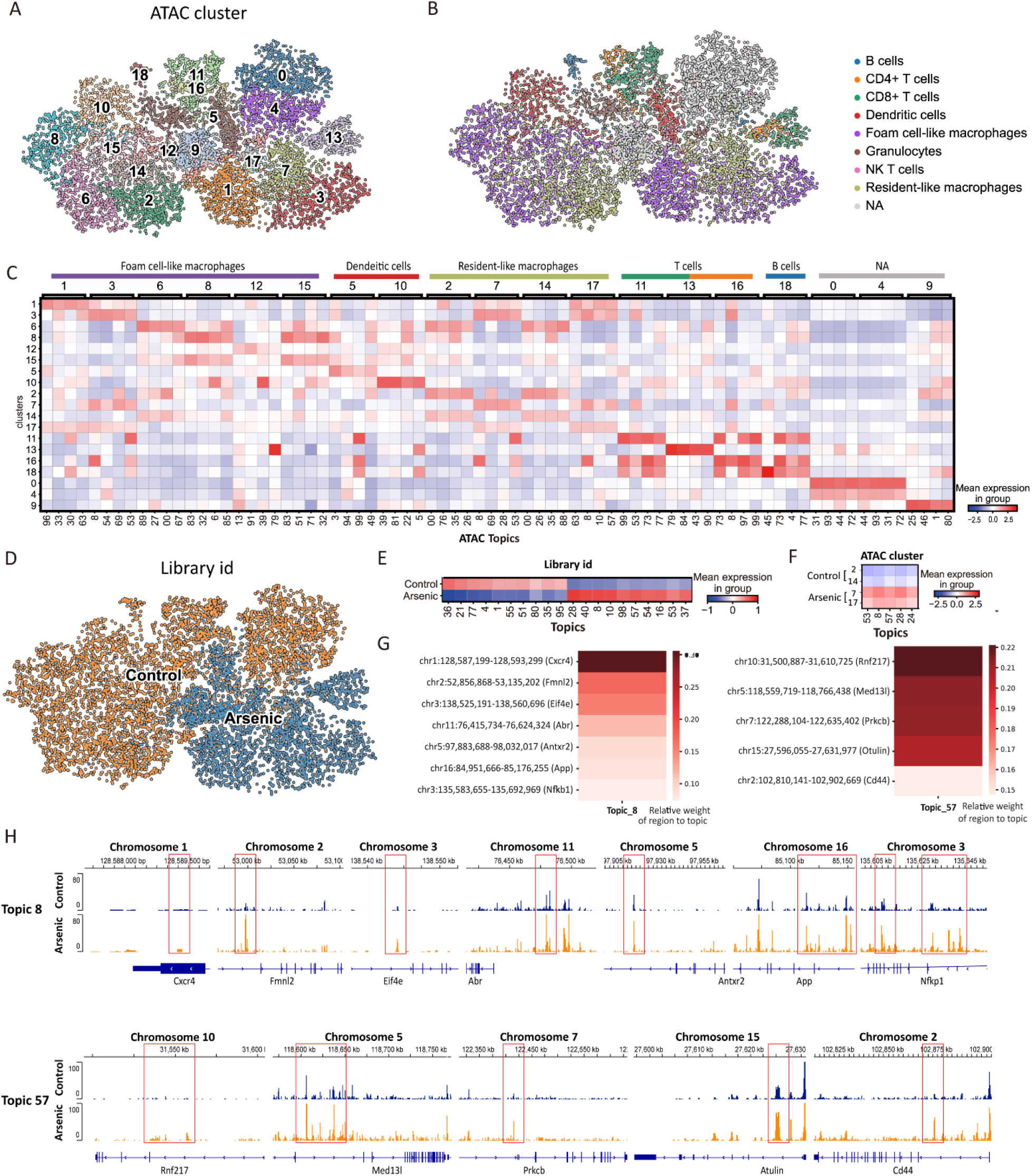
Exploring the scATAC-Seq dataset and its overlaps with that from scRNA-Seq. Uniform Manifold Approximation and Projection (UMAP) of the single cell gene expression data extracted from scATAC-Seq(m) performed on control and arsenic-exposed atherosclerotic plaques of apolipoprotein E knockout (apoE^-/-^) mice segregated into clusters using unsupervised clustering (A) and annotated for the immune cell types (B). Heatmap showing the topmost upregulated genes in each cluster defined in ‘B’ (C). Two dimensional UMAP of the gene expression data in single cells extracted from control (blue) and arsenic-exposed (orange) aorta of apolipoprotein E knockout (apoE^-/-^) mice from the scATAC-Seq(m) dataset (D). Heatmap showing the topics in control versus arsenic-treated samples (E). Heatmap showing the topics in control resident-like macrophage clusters 2 and 14 and arsenic-treated resident-like macrophage clusters 7 and 17 (F). Representative genes in topics 8 (left panel) and 57 (right panel) with the intensity of ‘red’ showing the relative weight of region to topic (G). Chromosomal view of the genes in topics 8 (top panel) and 57 (bottom panel) in control versus arsenic-treated samples (H).

To further examine the impact of arsenic on chromatin accessibility, we relabeled the UMAP based on the origin of single cells – plaques from control and arsenic-exposed animals (Figures 5D and S4). Interestingly, cells derived from control and arsenic samples occupied distinct areas on the UMAP based on their chromatin accessibility profiles. This underscores that arsenic-mediated alterations to the chromatin represent a key mechanism of arsenic toxicity and that these changes are analogous across different immune cell subtypes. We then examined the top differential topics between the control and arsenic-treated samples (Figure 5E). Topics 36, 21, and 77 were among the most representative in control samples, while topics 57, 28, and 8 were highly represented in cells derived from arsenic-treated samples.

To correlate the scATAC-Seq data with the two scRNA-Seq datasets, we examined the four resident-like macrophage clusters, clusters 2, 7, 14 and 17 in the scATAC-Seq analogous to clusters 5 and 6 in scRNA-Seq (m), and clusters 10 and 4 in the scRNA-Seq datasets (Figure 5F). Notably, clusters 2 and 14 exhibited topic signatures of control-derived samples, whereas clusters 7 and 17 were characteristic of arsenic-derived samples. We further examined topics 8 and 57 to reveal the encompassed genomic regions (Figure 5G), and visually represented the chromatin accessibility of some of these regions (enclosed in red boxes) along with their respective genes in control versus arsenic samples (Figure 5H). Genes *Cxcr4*, *Fmnl2*, *Eif4e*, *Abr*, *Antxr2*, *App* and *Nfkb1* are situated in the regions of interest in topic 8 and appear to exhibit increased chromosomal accessibility following arsenic exposure (Figure 5H). In topic 57, we discovered *Rnf217*, *Med13l*, *Prkcb*, *Otulin* and *Cd44* to show similar characteristics in the arsenic-treated group.

### In vitro validation of in vivo single cell datasets

Clusters 2, 14, 7 and 17 are comparable to the resident-like macrophage clusters 4 and 10 in sc-RNA-Seq and clusters 5 and 6 in scRNA-Seq (m). Therefore, in order to understand the arsenic mediated transcriptional changes, we explored the transcription factors differentially indicated in the CECs and AECs in the three single cell datasets (UMAP Figure 6A-C, transcription factors Figure 6D and Tables S13-S15 (scdiff2 results for scRNA-seq and scRNA-seq (m) and Tables S14-S15 for scATAC (m))). When compared, transcription factors IRF1, STAT1, YY1, POU2F1, CEBPA, CREB1, JUN and STAT6 (p-value: 1.13e-5) were differentially altered by arsenic in all three datasets (Figure 6E and Table S20). In addition to the substantial evidence sourced from scRNA-seq, scRNA-seq (m), and scATAC-seq (m) datasets, our study also incorporated the use of SDREM (38) as a potent analytical tool. SDREM helped bridge the gap between cell-cell interaction results and differentially expressed genes, and consequently revealed key transcription factors as pivotal nodes within the reconstructed signaling network. This additional layer of evidence underscores the value of SDREM in our analysis. SDREM’s distinct advantage lies in its ability to infer critical transcription factors from two dimensions. Firstly, it considers upstream receptors, identified through cell-cell interaction inference, using known protein-protein interactions. Secondly, it accounts for downstream target genes based on recognized TF-gene interactions. In the network, upstream receptors, depicted in blue, are considered the source nodes. The end target transcription factors responsible for the observed gene expression alterations are termed the network’s target nodes and are shown in green. Other proteins and transcription factors, which act as intermediaries in the signaling, are colored in orange.

**Figure 6.**
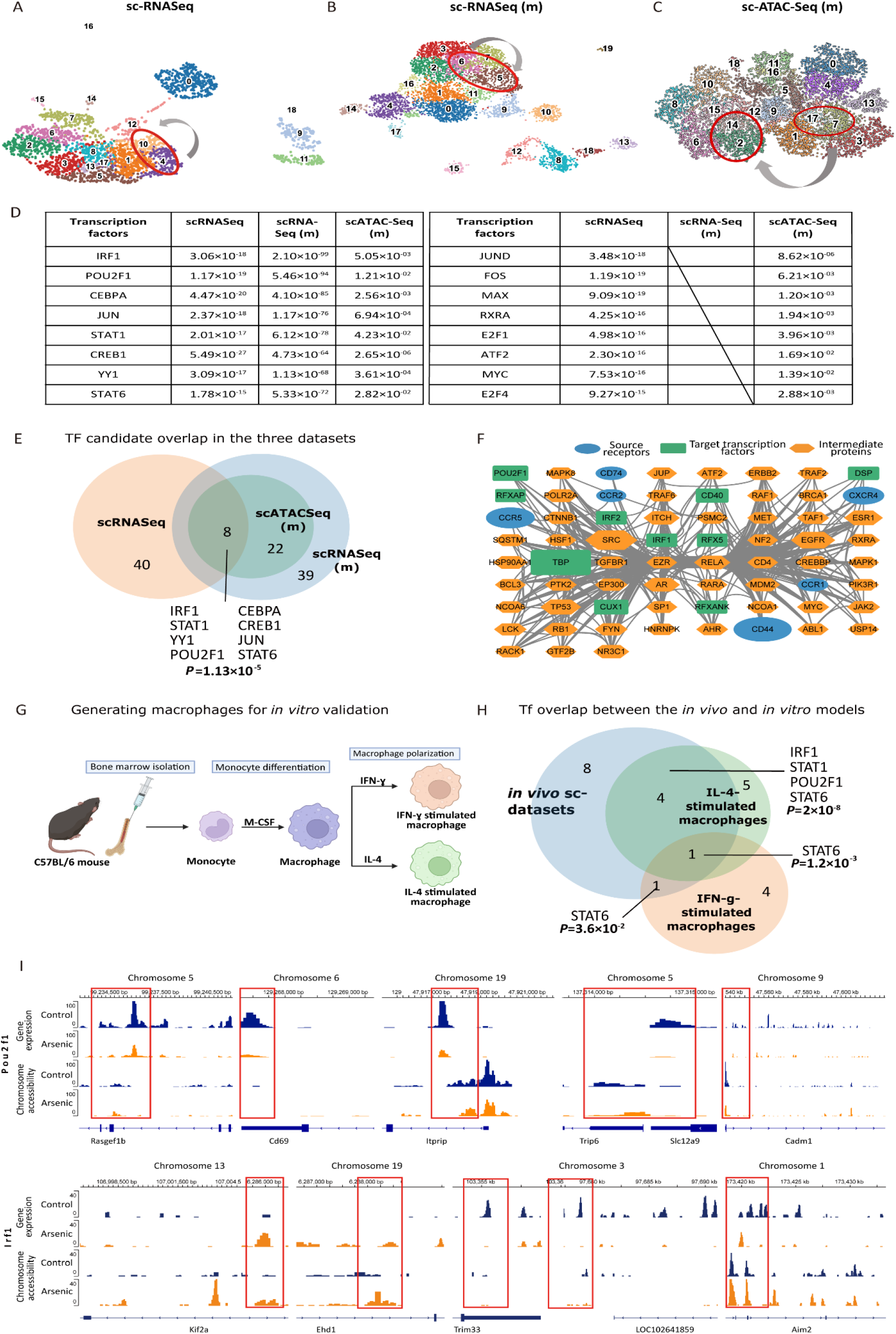
Comparative transcriptome analysis of mouse *in vivo* and *in vitro* datasets. UMAPs of scRNA-Seq, scRNA-Seq(m) and scATAC-Seq(m) datasets with resident-like macrophage clusters pursued in this study circled in red (A, B and C respectively). Table showing differential transcription factors between control and arsenic-treated samples in the aforementioned datasets with their p-values in the respective datasets (D). Wenn diagram showing transcription factors overlapping in each of the three datasets (E). Candidate transcription factors from the single cell datasets and a network of predicted receptors and intermediary proteins (F). Schematic for generating macrophages for in vitro validation of the in vivo single cell studies (G). Wenn diagram showing the overlap of the 8 transcription factors common to the single cell datasets and that in the in vitro-generated IFN-ɣ- and IL-4-stimulated macrophages along with their p-values (H). Chromosome accessibility and gene expression patterns for the genes under the transcriptional regulation of POU2F1 (top panel) and IRF1 (bottom panel) in the control and arsenic-treated resident-like cells in the single cell multiomic dataset (scRNA-Seq (m) and scATAC-Seq (m)) (I).

This dual perspective approach adds depth and validity to our study. The SDREM analysis notably highlights the significance of transcription factors IRF1 and POU2F1 in the regulation of distinct resident macrophage subtypes. To amplify the impact of these findings, we showcase the genomic regions regulated by these transcription factors, along with the differential transcriptomic and epigenomic data, to illustrate the effects of arsenic (Figure 6F).

To further validate these changes, we opted for in vitro models of macrophages. Atherosclerotic plaques are complex microenvironments and recapitulating them in vitro is challenging. Therefore, we utilized data from our previously published bulk RNA-Seq comparison of IFN-ɣ or IL-4-activated bone marrow-derived macrophages with or without concomitant arsenic exposure (Figure 6G and Tables S16-S19) (20). Based on the expression of target genes, STAT6 is the only transcription factor significantly altered in IFN-ɣ -polarized macrophages (p-value: 3.63e-2), and the only transcription factor common between the two in vitro-acquired macrophages and the in vivo single cell sequencing datasets (p-value: 1.2e-3) (Figure 6H and Table S20). Importantly, in IL-4-activated macrophages, transcription factors POU2F1, IRF1, STAT1 and STAT6 (p-value: 2.0e-8) were differentially altered, i.e., a 50% overlap with our multi-omics datasets (Figure 6H and Table S20). This suggests that the IL-4-activated macrophages (M2) highly resemble the resident-like macrophages in the single cell datasets and have the potential to model the arsenic-induced transcriptional and epigenomic changes.

To further explore the effects of arsenic on the four transcription factors commonly seen in our single cell and in vitro datasets, we delved into the genes under their regulatory control (Figure 6I). We draw attention to the genomic loci regulated by POU2F1 where arsenic induces changes in chromosome accessibility. This impacts the regions flanking *Rasgef1b*, *Cd69*, *Itprip*, *Trip6*, *Slc12a9*, and *Cdm1* genes, leading to significant gene expression changes (Figure 6I – top panel). Similarly, within the IRF1-regulated genomic locus, we observe changes in chromatin accessibility, presumably due to arsenic exposure. This manifests in the genomic regions flanking *Kif2a*, *Ehd1*, *Trim33*, and *Aim2*, yielding consequential gene expression changes (Figure 6I – bottom panel). However, data for the LOC102641859/*Rsf1* locus is limited, and hence a comprehensive assessment is challenging (Figure 6I – bottom panel).

### Overlap with human data

Several studies have sequenced atherosclerotic plaques from human subjects at a single cell level to decipher the resident cell types and their potential for targeted therapies. One such study integrated CyTOF, CITE-Seq and scRNA-Seq data from carotid artery plaques of patients undergoing carotid enderectomy to understand plaque microenvironment changes associated with adverse cardiovascular outcomes secondary to these plaques (39). As a result, this study identified macrophage clusters/gene signatures associated with patients that present with stroke or transient ischemic attack (symptomatic plaques) and those associated with no cerebrovascular event (asymptomatic plaques). Therefore, to evaluate the translational relevance of our murine model to the clinic, we compared the macrophage populations, their gene signatures, and their profiles in control versus exposed/treated subjects in our scRNASeq dataset to that in this clinical study (Figure 7A-F). Next, we focused on the resident-like macrophages of interest in our dataset and investigated potential analogies in human dataset. Interestingly, the gene signature of macrophages in arsenic-enriched cluster 4 and the control-enriched cluster 10 in our dataset align with that of clusters 7 and 6, respectively, from the human data, where key genes are enriched in corresponding clusters (Figure 7G). Importantly, the cluster 7 is more represented in symptomatic plaques, whereas cluster 6 aligns with asymptomatic plaques, which suggests that the arsenic exposure can potentially give rise to lesions with high susceptibility to adverse future cerebrovascular events (Figure 7G).

**Figure 7.**
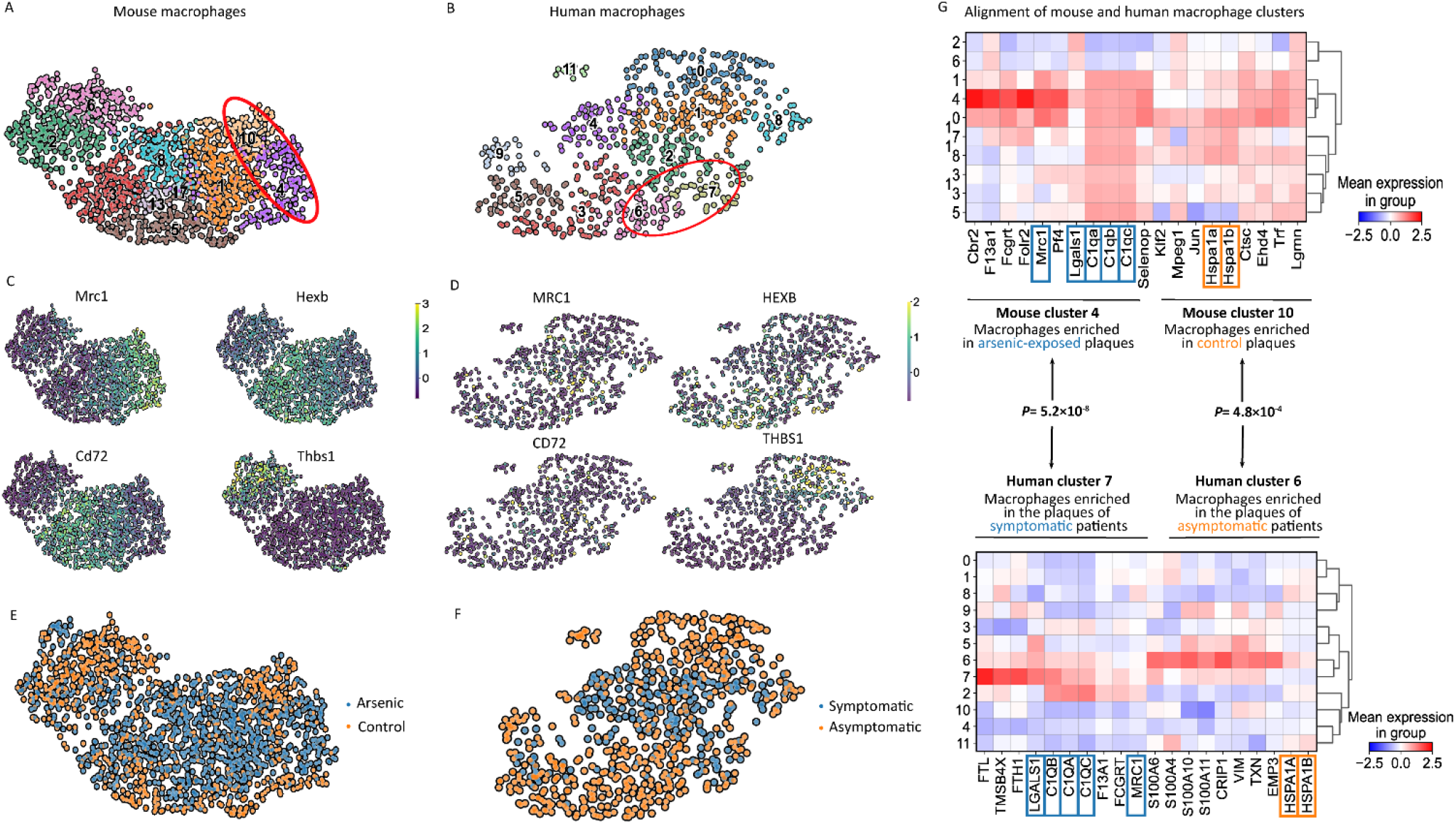
Comparative analysis of murine and human single cell datasets. UMAP of all macrophages from the scRNA-Seq dataset (A). UMAP of the human macrophages obtained from carotid artery plaques of patients undergoing carotid endarterectomy (B). Identification of macrophage subtypes using the previously determined gene signature for mouse (C) and human (D) datasets. UMAP of macrophages from mouse dataset segregated by control (orange) and arsenic-treated (blue) groups (E). UMAP of macrophages from human dataset segregated by their origin – plaques of asymptomatic (orange) and symptomatic (blue) patients (F). Alignment of the mouse resident-like macrophages 4 (arsenic-treated) and 10 (control) with those of human dataset 7 (symptomatic) and 6 (asymptomatic) (G).

## Discussion

To our knowledge, this is the first study that resolves atherosclerotic plaques at a single-cell level to understand environmental exposures, specifically the pro-atherogenic effects of arsenic. We investigated the diversity of CD45^+^ leukocytes populating the atherosclerotic plaques in male apoE^-/-^ mice on a high-fat diet and determined how arsenic altered this cellular milieu using scRNASeq and scATAC-seq. Our observed plaque cellular heterogeneity was consistent with previous reports (24, 25, 28, 39) and wasn’t drastically altered by arsenic. While the percentage of total plaque-resident macrophages remained consistent, arsenic specifically alters the relative abundance as well as the gene expression signatures in resident- and foam cell-like macrophages, driving potential changes in their cellular trajectories and interactions within the plaque microenvironment. Interestingly, our data suggest that arsenic affects the resident-like and foam cell-like macrophages in distinct fashions. Furthermore, arsenic significantly alters the epigenetic landscape of these plaques as implicated in our scATAC-Seq dataset, suggesting altered chromosomal accessibility as a potential mechanism of arsenic-mediated adverse atherosclerotic outcomes. Finally, when compared to a scRNASeq dataset from human plaques, arsenic-enriched resident macrophages corresponded to those of symptomatic rather than asymptomatic patients. Together, our data indicate that macrophages are key cellular targets of arsenic, and that arsenic changes the epigenetic/transcriptional landscape towards a phenotype characteristic of worse cardiovascular outcomes. This fits with the recent Scientific Statement from the American Heart Association defining metals as cardiovascular risk factors (40).

Arsenic is a known epigenetic modulator resulting in DNA methylation, histone modifications, and microRNA changes (41). For instance, Domingo-Relloso et al. reported a consortium of genomic positions differentially methylated by arsenic exposure in human blood from the Strong Heart Study cohort that associated with cardiovascular disease and a proportion of these genes were also significantly altered in mouse model of atherosclerosis as a result of arsenic exposure (42). We discovered that several genes in the differentially altered cistopics, such as Col1a1, Tgfbr1, and Nav2, etc., in our scATAC-Seq data corresponded to differentially methylated genes that correlated with arsenic exposure in this human dataset. Chromatin accessibility is strongly linked to DNA methylation. Increased DNA methylation leads to chromatin compaction and reduced accessibility for transcription factors, hindering their binding and impacting gene regulation (43–45). Consequently, low-methylated regions are typically observed in the promoters and enhancers of active genes, allowing for transcriptional activity. We show such changes in chromosome accessibility in the target genes of transcriptional factors such as IRF1 and POU2F1, which may alter the binding of these transcription factors in resident macrophages. Interferon Regulatory Factor 1 (IRF1) is an essential component of the interferon signaling cascade and an important modulator of macrophage function (46). While POU2F1 is not directly related to atherosclerosis, there is evidence of its correlation with myocardial fibrosis, a corollary of atherosclerotic ischemic disease, and with diabetes, a metabolic disease closely associated with atherosclerosis (47–50).

Intriguingly, the increased granularity provided by scRNASeq demonstrated that arsenic alters the transcriptome of the resident-like and foam-like macrophages uniquely. Supporting this conclusion, previous in vitro data from our lab demonstrated that arsenic resulted in distinct gene expression changes depending upon whether IFN-ɣ or IL-4 was used to polarize macrophages (20). This is intriguing as macrophages typically respond to their microenvironment and adopt a polarized functional state to respond to stimuli (51). However, addition of arsenic does not cause the same gene expression changes in resident versus foam-like macrophages, but in fact, results in opposite regulation of several important pathways (i.e., chemokine signaling).

The impact of arsenic at the transcriptome level was initially revealed through single-cell RNA-seq profiling in our study. However, the complexity of cellular state changes goes beyond the scope of a single modality, as even cells with similar transcriptomes can have dramatically different epigenomes. To delve deeper into the epigenetic shifts in immune cell populations upon arsenic exposure, we broke new ground by utilizing 10x multiome measurements. This added an additional layer of complexity to our unimodal single-cell RNA-seq analysis, enabling a more comprehensive characterization of these populations. The 10x multiome sequencing, which simultaneously profiles paired RNA-seq and ATAC-seq data from the same cell, offers an unrivaled perspective into the transcriptome and epigenome of the same cell set - a feat impossible with unimodal or unpaired multi-omics measurements. This dual analysis allows us to gain unprecedented insights into the effects of arsenic toxicity on various immune cell populations. To address our primary research objective—investigating the impact of arsenic toxicity on diverse immune cell populations—we strategically relied on the gene expression of classical markers for annotating the cell populations. This approach was favored over multimodal clustering of cells, which relies on learned joint cell embeddings inferred from multimodal data integration methods such as scGlue (52). The latter presents a potential drawback: lack of biological interpretability. This limitation could restrict our examination of immune cell populations as, for instance, two cell populations with similar epigenomes but divergent transcriptomes—and hence different cell types—could be misgrouped together. This outcome would result in cell clusters that do not align with established immune cell populations, which are typically characterized based on transcriptome and biomarker expression. In contrast, we prioritized annotating immune cell populations based on well-established biomarkers at the transcriptome level. We then leveraged scATAC-seq(m) measurements to investigate shifts in the epigenetic landscape upon arsenic exposure. Our compelling results from the scATAC-Seq (m) lead us to hypothesize that epigenetic regulation might be a pivotal mechanism driving the differential responses observed among macrophage types. As various stimuli are known to influence the polarization of macrophages through epigenetic changes (as reviewed in (53)), we suggest that arsenic may differentially affect macrophage subtypes due to their unique pre-programmed chromatin conformations. Alternatively, arsenic could modulate chromatin accessibility and gene expression during polarization or differentiation from monocytes. Future studies incorporating lineage tracing or time course experiments are recommended to further explore these possibilities, underscoring the utility of our chosen methodological approach.

Our study shows that arsenic has a profound effect on macrophage chromatin accessibility and gene expression in a murine model of atherosclerosis. However, it is important to note the caveats of our study. We assessed cellular heterogeneity of immune cells in arsenic-exposed apoE^-/-^ male mice on a high fat diet. Variables such as sex, diet, time of exposure, location of lesion, sample collection (i.e., preparation and sorting), and the loss of susceptible cell types during processing are important considerations in future experimental designs (54, 55). Plaque resident macrophage populations include inflammatory and interferon-inducible macrophages, but neither of these populations were resolved in our study. We enriched immune cell populations at the expense of other prominent cell types, such as endothelial and smooth muscle cells. There is considerable communication between these cells and immune cells, which the current experimental design may lack. Moreover, it is important to note that while single cell sequencing is a powerful tool to resolve the different cell types present in a tissue, it is limiting in providing spatial information. Nonetheless, this study provides a starting point to overlay the gene expression findings with immunohistochemistry studies and ask topological questions.

This study provides a framework to uncover the mechanisms of arsenic-enhanced atherosclerosis at the multi-omics, single-cell level, which may facilitate exploration of therapeutic approaches and novel markers. For example, our study suggests that arsenic exposure results in a profound change in epigenetics/chromatin accessibility, possibly due to DNA methylation changes. Folic acid supplementation, which increases the methyl donor S-adenosyl methionine, correlates with a decrease in the adverse effects caused by arsenic (56–60), potentially by counteracting arsenic-induced global hypomethylation or by increasing arsenic biotransformation through methylation (61, 62). Our findings, therefore, support testing folate supplementation as a means of mitigating arsenic-induced atherosclerosis, thus, providing potential public health solutions through single-cell multi-omics and precision toxicology (63).

## Supporting information

Supplementary Figs. S1 to S4 & Legends for Supplemetary Tables S1 to S20

## Acknowledgments

The authors acknowledge Yvhans Chery, Kathy Forner and Christian Young from the Lady Davis Institute of Medicine Research for their assistance in the initial stages of the experiment: animal handling and sample collection.

## Sources of Funding

Fonds de Reserche du Québec - Santé (KM)

Canadian Institutes of Health Research (CIHR) (PJT-180505 to JD; PJT-166142 to KKM)

Heart and Stroke Foundation of Canada (G-23-0034943 to KKM and JD)

Fonds de recherche du Québec - Santé (FRQS) (295298 to J.D., 295299 to JD)

Natural Sciences and Engineering Research Council of Canada (NSERC) (RGPIN2022-04399 to JD)

Meakins-Christie Chair in Respiratory Research (JD)

National Natural Science Foundation of China (62272278 & 61972322 to HW)

National Key Research and Development Program (2021YFF0704103 to HW)

Fundamental Research Funds of Shandong University (HW)

## Author Contributions

Conceptualization: KM, JD, KKM

Methodology: XH, KM, NS, NG, FD, TE, JD, HW, KKM

Investigation: KM, XY, JD, KKM

Visualization: XH, KM, JD, KKM

Supervision: HW, JD, KKM

Writing—original draft: KM, XY, JD, KKM

Writing—review & editing: KM, XY, JD, KKM

## Data and Materials Availability

All data needed to evaluate the conclusions in the paper are present in the paper and/or the Supplementary Materials. The raw data from scRNA-seq and multi-omics experiments and the processed data have been uploaded to GEO with the accession number GSE240753. The source code for process and analysis has been deposited in Zenodo @ https://doi.org/10.5281/zenodo.8286739 and in GitHub @https://github.com/mcgilldinglab/Arsenic_Toxicity_Single-Cell_Multi-Omics_Profiling. The visualization website results of scdiff2 have been deposited in Zen<u>odo @</u> https://doi.org/10.5281/zenodo.8305486.

