## Supplementary Figs. S1 to S4 & Legends for Supplemetary Tables S1 to S20 for "Unveiling the Impact of Arsenic Toxicity on Immune Cells in Atherosclerotic Plaques: Insights from Single-Cell Multi-Omics Profiling"

Kiran Makhani *et al.*

**This PDF file includes:**

Figs. S1 to S4

Legends for tables S1 to S20

**Other Supplementary Materials for this manuscript include the following:**

Tables S1 to S20

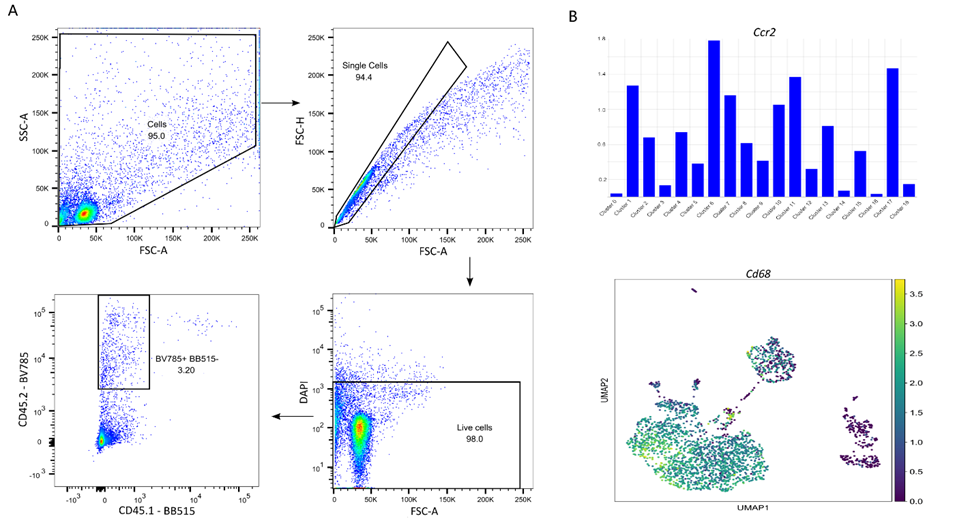

**Fig S1. Gating strategy and characterization**

Flow sorting gating strategy to obtain live CD45+ plaque resident cells for single cell sequencing (A). Relative expression of *Ccr2* (top) and *Cd68* (bottom) to characterize monocyte clusters (B).

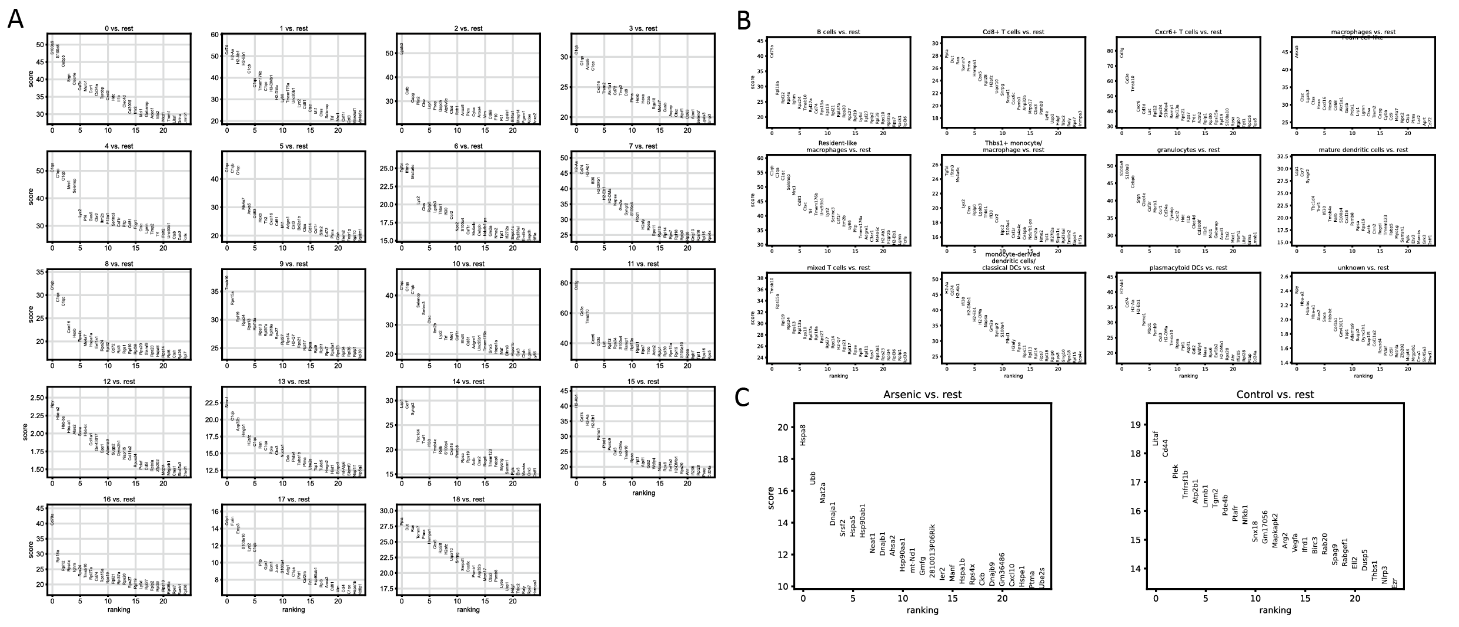

**Fig S2. Relative gene expression from scRNA-Seq dataset**

Gene signature of different immune cells and their expression profile relative to the rest of the immune cells as obtained from the scRNA-Seq dataset (A). Gene expression profile of each unsupervised cluster relative to rest of the clusters as obtained from the scRNA-Seq dataset (B). Differential gene expression profile of arsenic-enriched and control enriched clusters from scRNA-Seq dataset (C).

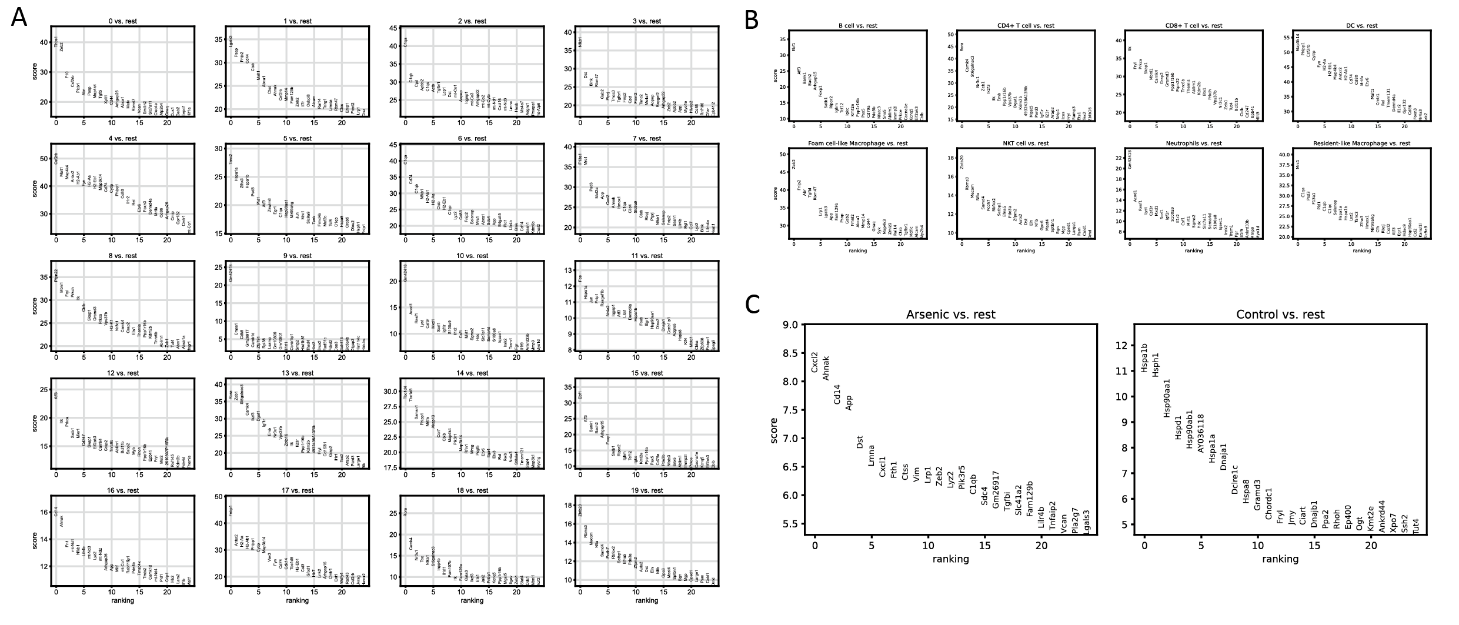

**Fig S3. Relative gene expression from sc RNA-Seq(m) dataset**

Gene signature of different immune cells and their expression profile relative to the rest of the immune cells as obtained from the scRNA-Seq(m) dataset (A). Gene expression profile of each unsupervised cluster relative to rest of the clusters as obtained from the scRNA-Seq(m) dataset (B). Differential gene expression profile of arsenic-enriched and control enriched clusters from scRNA-Seq dataset(m) (C).

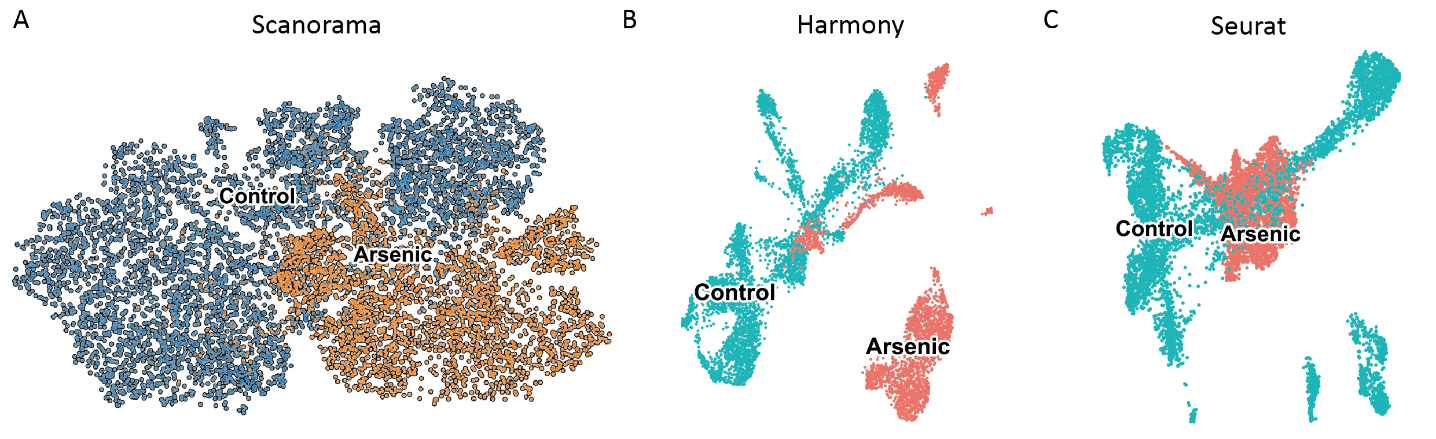

**Fig S4. Comparative analysis of integration methods for scATAC-Seq dataset**

UMAP showing integration of scATAC-Seq dataset using Scanoram (A). UMAP showing integration of scATAC-Seq dataset using Seurat (B). UMAP showing integration of scATAC-Seq dataset using Harmony (C).

**Table S1.**

Cell number of each cluster in scRNA-seq data.

**Table S2.**

Differential genes of each cluster in scRNA-seq data.

**Table S3.**

Enriched states of each cluster in scRNA-seq data.

**Table S4.**

Differential genes of cluster 4 v.s. cluster 10 in scRNA-seq data.

**Table S5.**

Differential genes of cluster 3 v.s. cluster 2 in scRNA-seq data.

**Table S6.**

Pathways of cluster 4 in scRNA-seq data.

**Table S7.**

Pathways of cluster 10 in scRNA-seq data.

**Table S8.**

Pathways of cluster 3 in scRNA-seq data.

**Table S9.**

Pathways of cluster 2 in scRNA-seq data.

**Table S10.**

Cell number of each cluster in scRNA-seq (m) data.

**Table S11.**

Differential genes of each cluster in scRNA-seq (m) data.

**Table S12.**

Significant status of each cluster in scRNA-seq (m) data.

**Table S13.**

Target genes of each candidate tf.

**Table S14.**

Common genes between genes from detopiced scATAC-seq data and target genes of each candidate tf.

**Table S15.**

Significant status of each candidate tf with genes from detopiced scATAC-seq data.

**Table S16.**

Common genes between M1 marker genes and target genes of each candidate tf.

**Table S17.**

Common genes between M2 marker genes and target genes of each candidate tf.

**Table S18.**

Significant status of each candidate tf with M1 marker genes.

**Table S19.**

Significant status of each candidate tf with M2 marker genes.

**Table S20.**

Overall common tf candidates status.
